# Plakoglobin is a mechanoresponsive regulator of naïve pluripotency

**DOI:** 10.1101/2022.03.13.484158

**Authors:** Timo N. Kohler, Joachim De Jonghe, Anna L. Ellerman, Ayaka Yanagida, Michael Herger, Erin M. Slatery, Katrin Fischer, Carla Mulas, Alex Winkel, Connor Ross, Sophie Bergmann, Kristian Franze, Kevin Chalut, Jennifer Nichols, Thorsten E. Boroviak, Florian Hollfelder

## Abstract

Biomechanical cues are instrumental in guiding embryonic development and cell differentiation. Understanding how these physical stimuli translate into transcriptional programs could provide insight into mechanisms underlying mammalian pre-implantation development. Here, we explore this by exerting microenvironmental control over mouse embryonic stem cells (ESCs). Microfluidic encapsulation of ESCs in agarose microgels stabilized the naïve pluripotency network and specifically induced expression of Plakoglobin (*Jup*), a vertebrate homologue of β-catenin. Indeed, overexpression of Plakoglobin was sufficient to fully re-establish the naïve pluripotency gene regulatory network under metastable pluripotency conditions, as confirmed by single-cell transcriptome profiling. Finally, we found that in the epiblast, Plakoglobin was exclusively expressed at the blastocyst stage in human and mouse embryos – further strengthening the link between Plakoglobin and naïve pluripotency *in vivo*. Our work reveals Plakoglobin as a mechanosensitive regulator of naïve pluripotency and provides a paradigm to interrogate the effects of volumetric confinement on cell-fate transitions.

**Highlights:**

- 3D agarose spheres stabilize the naïve pluripotency network in mouse ESCs.
- Volumetric confinement induces expression of Plakoglobin, a vertebrate homologue of β-catenin.
- Plakoglobin expression in the epiblast is specific to pre-implantation human and mouse embryos.
- Plakoglobin overexpression maintains naïve pluripotency independently of β-catenin.

## Introduction

Cells integrate cytokine-mediated, chemical and physical stimuli from the extracellular environment to regulate cell fate and function. Even the physical properties of a cell itself, including cortical stiffness^1^, membrane tension^2, 3^ and intracellular crowding^4^, are subject to dynamic changes in response to external cues, thereby integrating microenvironmental information into a physiological response^5, 6^. This intricate pattern of regulation is of particular importance during embryogenesis, when pluripotent cells are faced with the complex task of undergoing spatiotemporal lineage specification to generate an entire fetus.

The initial state of this process, naïve pluripotency, emerges in the epiblast of the mouse blastocyst prior to implantation and can be captured *in vitro* as mouse embryonic stem cells (ESCs)^7, 8^. ESCs correspond to the pre-implantation epiblast^9^ and retain full developmental capacity^10–12^ when injected into a host embryo^11, 13^. This naïve ‘ground state’ of pluripotency can be sustained indefinitely in a defined culture regime termed 2i/LIF, where self-renewal is mediated through inhibition of mitogen activated kinase kinase (MEK) (by PD0325901, PD) and glycogen synthase kinase-3 (GSK-3β) (by CHIR99021, CH), as well as stimulation of JAK/STAT signaling by leukemia inhibitory factor(LIF)^12^. Inhibition of the MEK/ERK cascade suppresses developmental progression towards post-implantation stages^14, 15^, GSK-3 inhibition stabilizes β-catenin, which supports the naïve pluripotency circuitry through de-repression of TCF7L1 (also TCF3)^16, 17^ and LIF stimulates transcription of key pluripotency factors *Klf4* and *Tfcp2l1*^18, 19^. Any two of the three 2i/LIF components (PD, CHIR and LIF) are sufficient to maintain naïve pluripotency^20^. In addition to chemical and cytokine-mediated signals, biomechanical cues are emerging key players in the regulation of embryonic^21–25^ and adult stem cells^4, 26–28^. However, the identity of mechanoresponsive genes in ESCs and the mechanisms transforming physical responses into transcriptional programs remain poorly understood.

ESCs become sensitized towards mechanical stress^29^ and decrease membrane tension upon exit from naïve pluripotency^3^. Membrane tension regulates the endocytic uptake of MEK/ERK signaling components and thus somatic cell-fate acquisition^2^. Recently, volumetric compression has been identified as a physical cue to promote self-renewal of intestinal stem cells in organoid cultures^4^, and dedifferentiation of adipocytes^30^, via WNT/β-catenin signaling. Volumetric compression induces molecular crowding, which stabilizes LRP6 signalosome formation and consequently elevates WNT/β-catenin signaling. In the blastocyst, hydrostatic pressure also plays a critical role in embryo size regulation and cell fate specification^31, 32^. However, little is known about the effects of volumetric compression and spatial confinement in naïve pluripotent ESCs.

Considering the powerful effect of WNT/β-catenin signaling on naïve pluripotency^16, 33^ and its link to molecular crowding, we sought to systematically interrogate volumetric confinement in ESCs. Biomimetic culture systems that emulate volumetric compression in the developing blastocyst may provide a route to analyzing developmental processes under conditions of a crowded native environment in the embryonic niche. Here, we generated spherical agarose scaffolds^34–36^ to encapsulate naïve pluripotent ESCs in microgels and analyze their transcriptional and morphological development. We identify Plakoglobin (*Jup*), a vertebrate homologue of β-catenin, as a mechanoresponsive gene and potent regulator of naïve pluripotency.

## Results

### Microgel culture of embryonic stem cells stabilizes the pluripotent state

To investigate the effects of 3D-culture on pluripotency we encapsulated mouse ESCs into agarose microgels (**Figure 1A, 1B**). This was performed in microfluidic flow-focusing chips by generating monodisperse water-in-oil emulsion droplets containing liquid low-melting agarose (at 37 °C) that, upon cooling on ice, formed biologically inert, spherical 3D hydrogel scaffolds (**Figure S1A-B**). The resulting cell-containing microgels were cultured under self-renewing (naïve pluripotent: 2i/LIF; metastable pluripotent: serum/LIF) or differentiating (N2B27) conditions as suspension culture (**Figure 1C, S1C-D**). Encapsulated ESCs in 2i/LIF displayed homogeneous expression of the general pluripotency marker OCT4 and the naïve pluripotency marker KLF4. These were downregulated and absent respectively, when cultured in N2B27 for 48 h (**Figure 1C**). When metastable pluripotent (serum/LIF) ESCs were cultured in microgels the cells formed tightly packed colonies with indistinguishable cell boundaries and acquired a round, dome-shaped morphology, similar to that seen in naïve cells (2i/LIF) and unlike the serum/LIF cells cultured on plastic (**Figure 1B, S1C**). To determine the effects of 3D microgel culture on pluripotent ESCs, we utilized the *Rex1::GFPd2* reporter cell line (RGd2) expressing a destabilized GFP from the endogenous REX1 (gene name *Zfp42*) locus as a quantitative real-time readout for naïve pluripotency^16^ (**Figure 1D**). In this system, loss of Rex1-GFPd2 indicates exit from naïve pluripotency^16, 33^ (**Figure 1D, S1E**). Consistent with previous reports^37^, in 2D culture, 2i/LIF induced uniformly high (∼99% of cells) reporter expression, in contrast to serum/LIF, which had a bimodal distribution (**Figure 1E**). Encapsulated ESCs in 2i/LIF showed homogenous (∼99% of cells) Rex1-GFPd2 levels, similar to conventional ESCs in 2i/LIF. However, microgel suspension culture in serum/LIF revealed a ∼20% increase in Rex1-GFPd2 expression compared to serum/LIF on tissue culture plastic (**Figure 1F**). This observation is mirrored by other naïve pluripotency markers: we stained for KLF4 and found a robust increase in KLF4 protein levels in serum/LIF microgel-cultured ESCs over tissue culture plastic (**Figure 1G**). Next, we tested the developmental potential of encapsulated ESCs in chimera assays. H2B-tq labelled ESCs were cultured for 48 h in agarose microgel suspension culture, released from the gels by agarase treatment and microinjected (3-5 cells/embryo) into 8-cell stage host embryos (n = 11). Microgel-cultured ESCs robustly colonized the pluripotent epiblast compartment, as indicated by colocalized SOX2 (epiblast marker) immunofluorescence staining, with an efficiency of more than 70% (**Figure 1H**). We conclude that 3D agarose microgel culture is suitable for the culture of naïve pluripotent (2i/LIF) ESCs and stabilizes pluripotency under metastable pluripotent (serum/LIF) culture conditions.

**Fig. 1.**
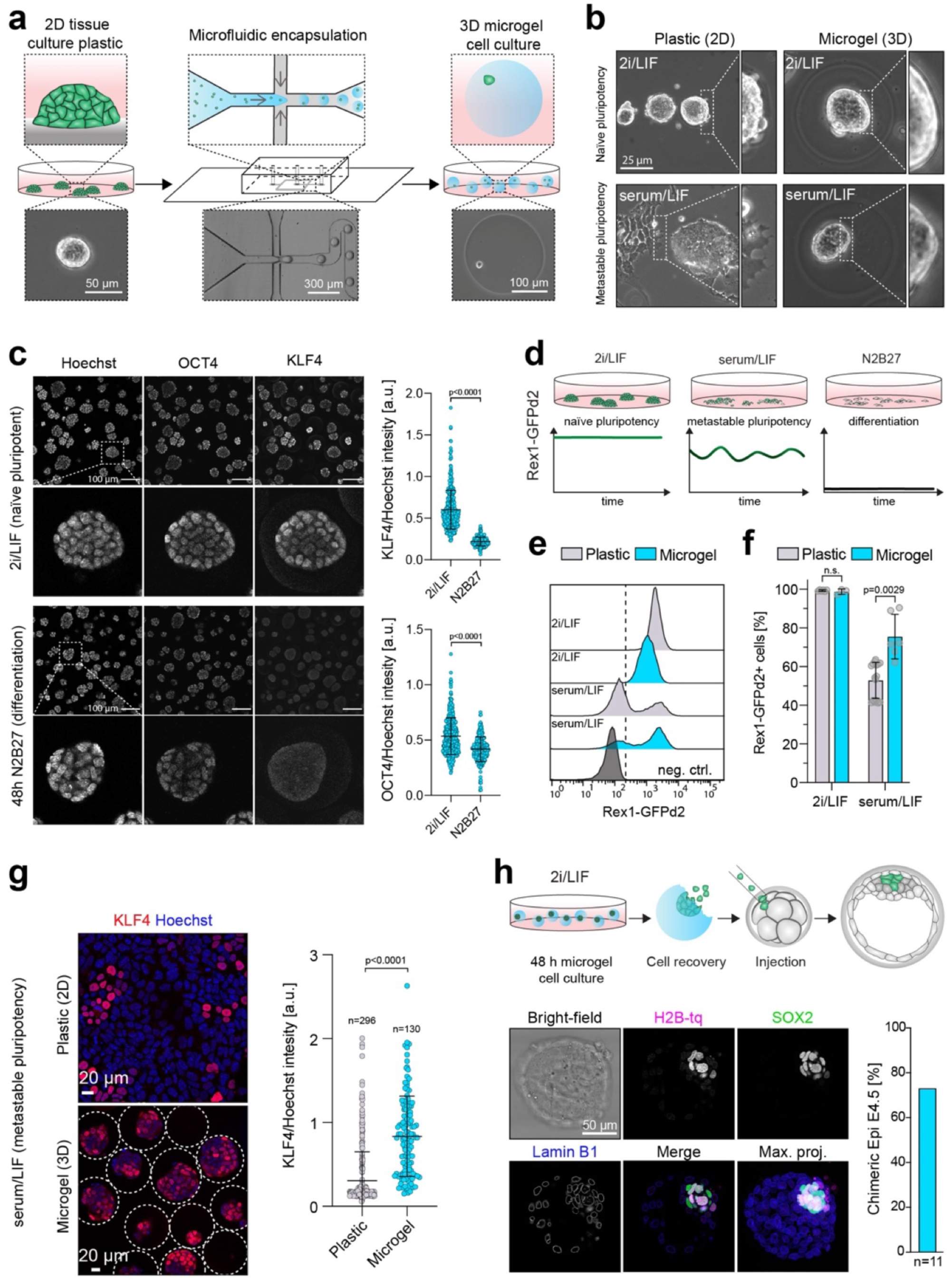
Agarose microgel encapsulation of mouse ESCs supports naïve pluripotency. **a**, Schematic illustration of the microfluidic cell encapsulation process. Tissue culture plastic (2D) mouse ESCs were single cell separated and microfluidically compartmentalized into agarose-in-oil microdroplets. Upon polymerization and de-emulsification, spherical cell-laden agarose microgels (3D) were cultured in suspension culture. **b**, Phase contrast images of ESC cultured on tissue culture plastic or encapsulated in microgels in naïve pluripotent (2i/LIF) or metastable pluripotent (serum/LIF) conditions. Microgel cultured serum/LIF cultured cells acquire a naïve-like phenotype. **c**, Confocal immunofluorescence images of microgel encapsulated ESCs stained for the general pluripotency marker OCT4 and the naïve pluripotency marker KLF4 after being cultured for 48 h in 2i/LIF (naïve pluripotent) or N2B27 (differentiation). KLF4 and OCT4 intensities (as shown on the right) were normalized to Hoechst. Error bars indicate the mean and standard deviations. n = 400 (2i/LIF), n = 285 (N2B27). **d**, The *Rex1::GFPd2* (RGd2) reporter system allows near real-time (destabilized GFP with a half-life of 2 h) analysis of pluripotency. Homogenous expression (naïve in 2i/LIF), heterogeneous expression (metastable in serum/LIF) and loss of expression upon exit from pluripotency (N2B27). **e&f**, Flow cytometric analysis of RGd2 ESCs cultured on plastic *vs* in microgels. No differences were observed between cells cultured under naïve conditions (2i/LIF). An increase in Rex1-GFPd2+ cells under metastable conditions (serum/LIF) when cultured for 48 h in microgels suggests a shift towards naïve pluripotency. Negative control: wildtype ESCs. Error bars indicate the mean and standard deviations. N = 6 (plastic 2i/LIF), N = 3 (microgel 2i/LIF), N = 16 (plastic serum/LIF), N = 6 (microgel serum/LIF). **g**, Confocal immunofluorescence images of serum/LIF cultured (microgel and plastic) ESCs stained for KLF4. The increase in KLF4+ cells suggests a shift towards naïve pluripotency in microgel cultured cells. KLF4 intensities (as shown on the right) were normalized to Hoechst. Error bars indicate the mean and standard deviations. n = 296 (plastic), n = 130 (microgel). **h**, Injection of microgel-cultured naïve ESCs (nuclear H2B-tq reporter) into 8-cell stage embryos lead to ∼73% (n = 8/11) blastocysts chimeras 48 h after *in vitro* culture. Injected cells were identified via the H2B-tq reporter. Cells were stained for Lamin B1 (outlines all nuclei) and SOX2 (Epiblast).

### Microgel suspension culture induces Plakoglobin expression

To delineate the microgel-induced changes in the transcriptional program of pluripotent cells, we performed RNA-seq of naïve ESCs cultured on tissue culture plastic *versus* microgel encapsulated naïve ESCs. Pearson correlation and principal component analysis (PCA) confirmed that samples clearly separated according to experimental conditions (**Figure S2A-B**). General pluripotency factors remained robustly expressed in both conditions, with the core pluripotency factors *Pou5f1* and *Sox2* being slightly elevated in 3D-microgel cultured cells (**Figure 2A**). Additionally, we observed a global increase across the expression of naïve pluripotency factors such as *Esrrb*, *Klf2*, *Klf4*, *Klf5*, *Tbx3* and *Tfcp2l1* in encapsulated ESCs (**Figure 2B**). Transcriptome-wide comparison showed that some of the most differentially expressed genes were the naïve pluripotency factors *Klf2* and *Klf4*, components of the WNT/β-catenin signaling pathway, including *Sfrp1* and *Esrrb* (naïve pluripotency associated transcription factor and downstream target of β-catenin) as well as the β-catenin vertebrate homologue *Jup* (Plakoglobin) (**Figure 2C**). We performed gene set enrichment analysis (GSEA) of 2D-tissue culture plastic *versus* encapsulated ESCs and obtained enrichment for “PluriNetWork”, “Regulation of Actin Cytoskeleton”, “Focal Adhesion” and “WNT Signaling” for ESCs cultured in agarose microgels (**Figure S2C-D)**. Considering the important role of the WNT/β-catenin signaling for self-renewal and pluripotency in the mouse^16^ and β-catenin’s crucial role in the formation of adherens junctions, we examined individual members of adherens junction-associated genes: *Ctnna1* (α-catenin), *Ctnnb1* (β-catenin), *Jup* (Plakoglobin / γ-catenin), *Ctnnd1* (p120 / δ-catenin), and *Cdh1* (E-cadherin). *Cdh1*, *Ctnna1* and *Ctnnb1* expression remained mostly unchanged, while *Jup* and *Ctnnd1* were significantly upregulated (**Figure 2D, S2E**). *Jup* showed the strongest (∼6-fold) increase in microgels. Immunofluorescence staining confirmed strong induction of Plakoglobin (*Jup*) upon agarose microgel culture at the protein level and demonstrated localization to adherens junctions (**Figure 2E, S2E**). β-catenin was expressed in ESCs cultured in both conditions, although we noted slightly higher cytoplasmic protein levels in 2D cultures on plastic (**Figure 2F, S2E**). These results demonstrate that microgel culture induces Plakoglobin, which localizes to adherens junctions at the expense of β-catenin.

**Fig. 2.**
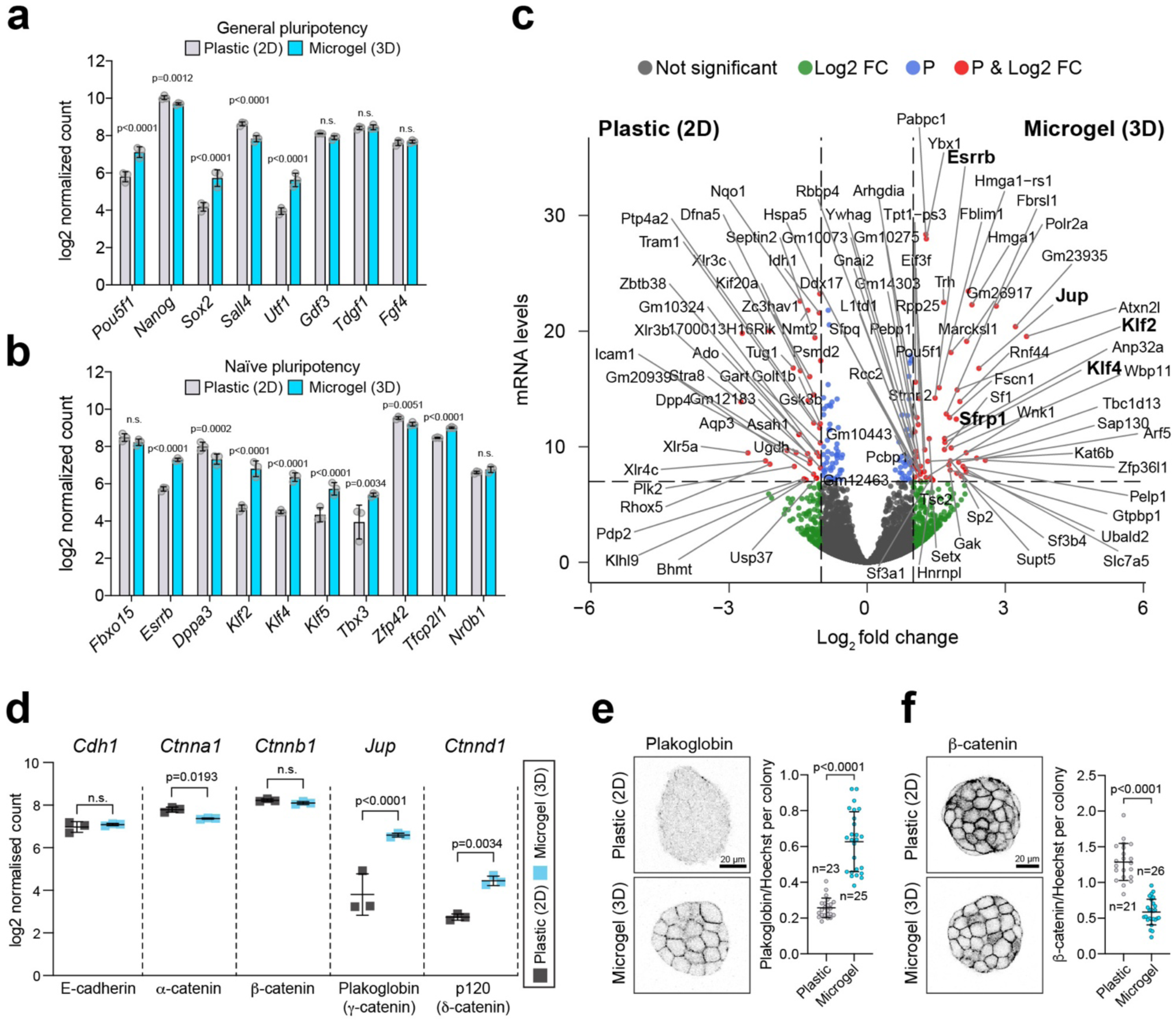
Transcriptomic analysis of encapsulated ESCs reveals upregulation of Plakoglobin. **a&b**, Bulk RNA-seq analysis of general (a) and naïve (b) pluripotency-associated genes for ESCs seeded either on plastic (gray) or encapsulated in microgels (blue) under 2i/LIF culture conditions. Error bar indicate the mean and standard deviations. **c**, Volcano plot representing the list of differentially expressed genes between ESCs cultured on tissue culture plastic (negative log2 fold changes in gene expression) or in agarose microgels (positive log2 fold changes in gene expression) in 2i/LIF. Log2 fold change cut-off was |log2 fold change| > 1 and Bonferroni-adjusted p-values cut-off was 10^-7. NS stands for not-significant (gray), Log2 FC (green color) indicates gene that passed the log2 fold change cut-off, P (blue) indicates genes that passed p-value cut-off, P&Log2 FC (red) indicates genes that passed log2 FC and p-value cut-offs. **d**, 3D microgel culture lead to a significant up-regulation at gene expression level of Plakoglobin (*Jup*) and p120 (*Ctnnd1*) compared to 2D plastic culture conditions. Error bars indicate the mean and standard deviations. **e**, Confocal immunofluorescence images of naïve pluripotent (2i/LIF) ESCs cultured on plastic and encapsulated in microgels stained for Plakoglobin. Plakoglobin levels were quantified for colonies gown on plastic (n = 23) and in microgels (n = 25). Error bars indicate the mean and standard deviations. **f**, Confocal immunofluorescence images of naïve pluripotent (2i/LIF) ESCs culture on plastic and encapsulated in microgels stained for β-catenin. β-catenin levels were quantified for colonies gown on plastic (n = 21) and in microgels (n = 26). Error bars indicate the mean and standard deviations.

### Plakoglobin expression is regulated via microgel-mediated volumetric confinement

To delineate microgel-induced volumetric confinement from general effects of 3D-culture, we performed time course analysis of ESC microgel culture side-by-side with hanging drop cultures (**Figure 3A**). Hanging drop culture allows ESCs to grow in suspension as 3D-aggregates, but in the absence of confinement from the microgel scaffold. Remarkably, under naïve conditions only microgel encapsulated cells, but not cells cultured in a hanging drop, showed an upregulation of Plakoglobin – an effect observed as early as 48 h after encapsulation (**Figure 3B, S3A**). In both conditions the naïve marker KLF4 remained strongly expressed (**Figure 3B**). To exclude potential aggregate size-induced differences, we started both culture regimes from single cells to obtain structures of the same size (**Figure 3C-D**). β-catenin was consistently detected in all conditions tested, localized predominantly towards the cell membrane and exhibited slightly elevated levels in microgels compared to hanging drops (**Figure 3C**). In contrast, Plakoglobin expression was exclusively detected in microgel-cultured cells but not in cells cultured in hanging drops (**Figure 3D**). In 2D culture and absence of microgel-mediated volumetric confinement, Plakoglobin levels remained low and were unaffected by matrix stiffness (**Figure S3B**). We further probed whether microgel confinement was required to maintain Plakoglobin expression by releasing ESCs after 48 h from microgels followed by 48 h of hanging drop suspension culture. β-catenin was expressed in all conditions (**Figure 3C**). However, Plakoglobin expression of encapsulated ESCs was lost upon release from gels and transfer to hanging drop cultures (**Figure 3D**). This result demonstrates that Plakoglobin upregulation and maintained expression is directly dependent on microgel-mediated volumetric confinement, but independent of 3D-culture.

**Fig. 3.**
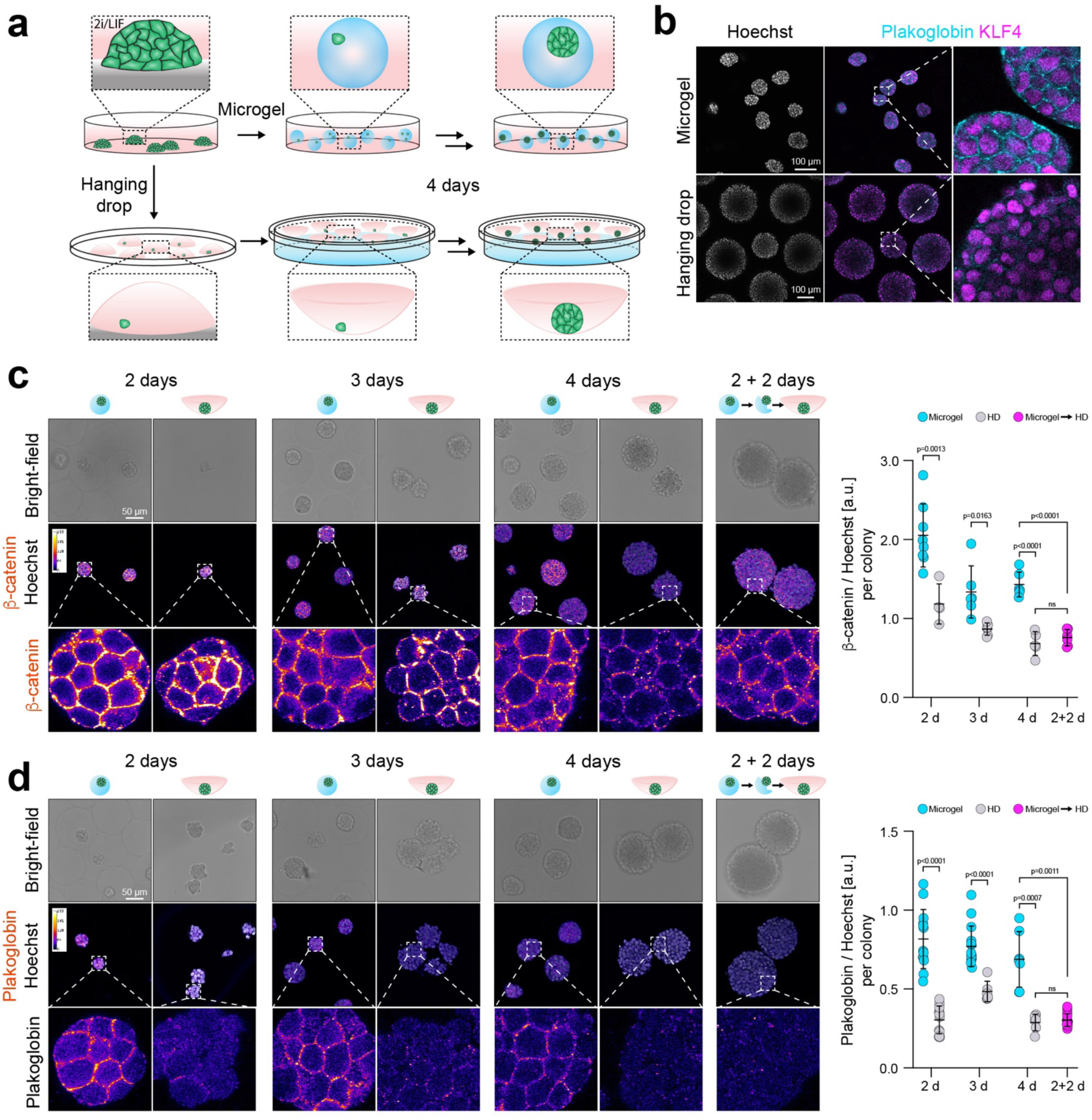
Microgel-mediated volumetric confinement induces mechanoresponsive Plakoglobin expression in ESCs. **a**, Schematic 3D cell culture formats: in microgel encapsulation providing a structural scaffold (top) vs in hanging drops without providing a scaffold (bottom). **b**, Confocal immunofluorescence images of naïve pluripotent (2i/LIF) ESCs cultured as hanging drops or microgel-encapsulated. Cells were stained for Plakoglobin (PG) and KLF4 after 48 h in culture. **c&d**, Confocal immunofluorescence images of single-cell cultured naïve pluripotent (2i/LIF) ESCs in hanging drops and microgels for 96 h. Cells were stained after 48 h, 72 h, 96 h and 48 h in microgel followed by 48 h in hanging drops (following release from the gels) for (c) β-catenin and (d) Plakoglobin. Plakoglobin expression was mostly detected in microgel cultured cells whereas β-catenin was expressed in hanging drop and microgel cultured cells. β-catenin and Plakoglobin levels were quantified per colony (shown on the right). Error bars indicate the mean and standard deviations.

### Plakoglobin expression is a defining feature of the naïve pre-implantation epiblast in mouse and human embryos

Considering that volumetric confinement induced Plakoglobin in microgel-encapsulated pluripotent ESCs, we sought to determine *Jup*/Plakoglobin expression in the early embryo – the origin of naïve pluripotency. *Jup* transcription was highest in the pluripotent compartment (E3.5-ICM and E4.5-EPI) of the pre-implantation mouse embryo (**Figure 4A, S4A**), consistent with *Jup* expression in the naïve pluripotent ESCs. Notably, *Jup* transcription was sustained in diapause, a facultative developmental arrest at the blastocyst stage (Ketchel et al., 1966; Scognamiglio et al., 2016), but downregulated in the post-implantation epiblast (**Figure 4A, S4A**). This stands in contrast to *Ctnnb1*, which was robustly expressed throughout pre-implantation development and further upregulated upon implantation (**Figure 4B, S4B**). At the protein level, we did not detect Plakoglobin at the 8-cell stage (E2.5), but obtained a robust membrane-adjacent signal of both the trophectoderm and the ICM at the blastocyst stage (E3.5-4.5), in which pluripotency is established (**Figure 4C**). Strikingly, Plakoglobin was absent in the primed pluripotent, epithelialized epiblast of the early post-implantation egg-cylinder at (E5.5-E6.5), while it remained in the extraembryonic ectoderm (ExE) (**Figure 4D**). In contrast, β-catenin did not exhibit stage-specific patterns and was localized cell membrane adjacent throughout all stages analyzed (**Figure 4E-F**), in agreement with the transcriptome data (**Figure 4B**). To investigate whether Plakoglobin expression was restricted to the pluripotent epiblast cells of the ICM, we performed immunofluorescence staining in hatched blastocysts at E4.25 for Plakoglobin, the epiblast marker OCT4 and the primitive endoderm (PrE) marker SOX17 (**Figure 4G**). Plakoglobin expression was detectable in SOX17 positive cells where the PrE had not sorted to the ICM surface and not yet formed a continuous epithelium. However, in slightly later stages when the PrE has formed a mature epithelium Plakoglobin expression was downregulated (**Figure 4G, S4B**). Due to the high degree of evolutionary conservation of β-catenin and Plakoglobin^38^, we investigated Plakoglobin’s expression pattern in human embryos for comparison. Immunofluorescence staining of Plakoglobin revealed its absence in early cleavage embryos at the 4-cell stage, in contrast to β-catenin (**Figure 4H**). However, consistent with mouse development, human embryos at the blastocyst stage, robustly displayed Plakoglobin signal in the trophectoderm and the pluripotent epiblast **(Figure 4I, S4C**). Co-staining of Plakoglobin with the epiblast markers SOX2 and the hypoblast marker SOX17 showed that hypoblast fate is linked to a reduction in Plakoglobin levels (**Figure 4I**). We conclude that Plakoglobin is an evolutionary conserved feature of the mouse and human pre-implantation pluripotent epiblast.

**Fig. 4.**
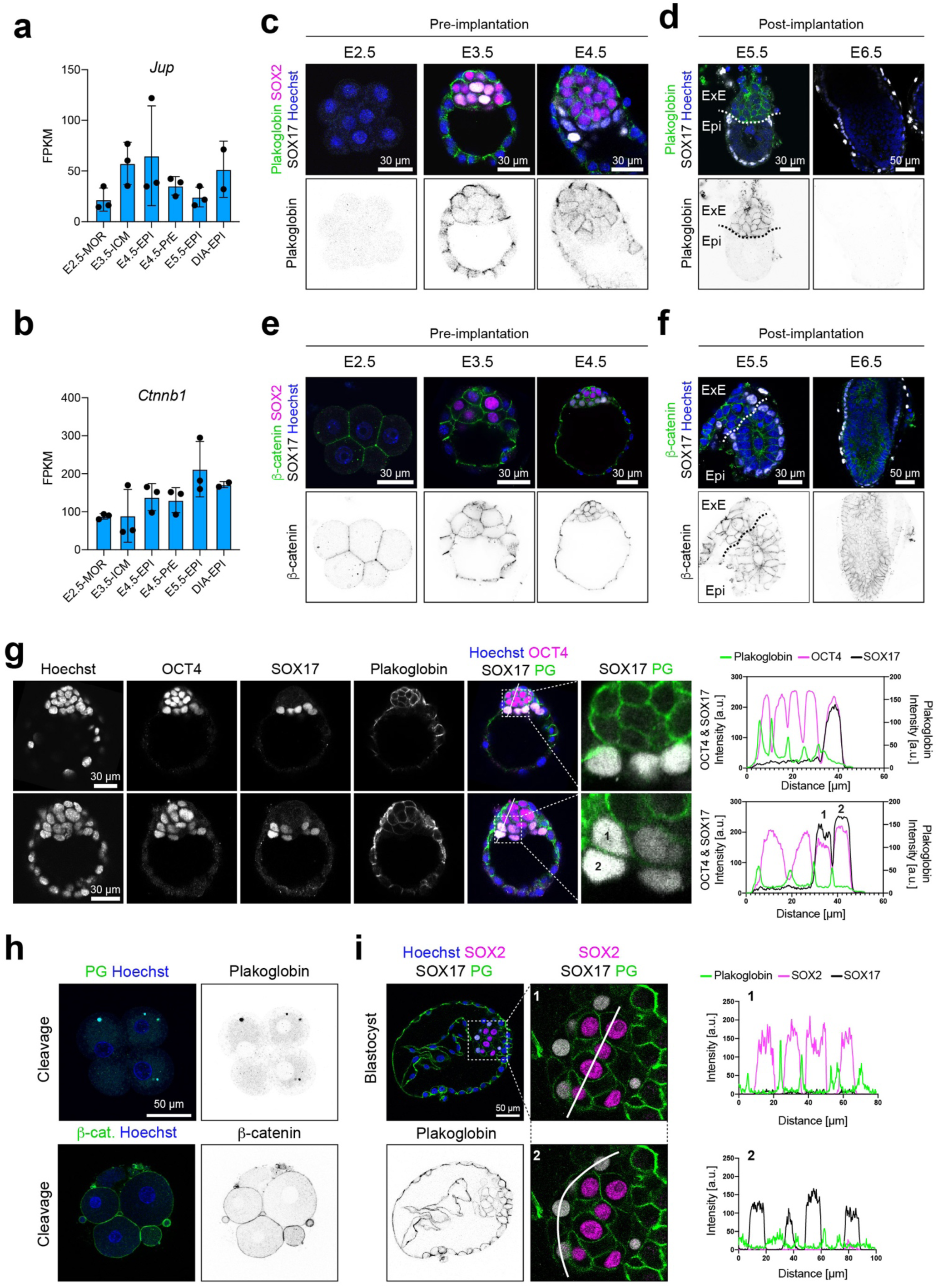
Plakoglobin expression is evolutionarily conserved and associated with pre-implantation pluripotency. **a**, Meta-analysis of RNA-seq data^10^ in Fragments Per Kilobase Million (FPKM) for *Jup* (Plakoglobin) during early embryogenesis. **b**, Meta-analysis of RNA-seq data^10^ in FPKM for *Ctnnb1* (β-catenin) during early embryogenesis. **c**, Confocal immunofluorescence images of pre-implantation embryos (E2.5, E3.5 and E4.5) stained for Plakoglobin, SOX2 (epiblast marker) and SOX17 (primitive endoderm marker, pre-implantation). Plakoglobin expression in the epiblast coincides with the emergence of naïve pluripotency in the pre-implantation epiblast. Scale bar: 30 µm. **d**, Confocal immunofluorescence images of post-implantation embryos (E5.5 and E6.5) stained for Plakoglobin and SOX17 (Visceral endoderm marker, post-implantation). In the post-implantation embryo Plakoglobin expression was restricted to the extraembryonic ectoderm (ExE) and completely absent in the epiblast (Epi) at E5.5. Scale bar: 30 µm. **e**, Confocal immunofluorescence images of pre-implantation embryos (E2.5, E3.5 and E4.5) stained for β-catenin, SOX2 (epiblast marker) and SOX17 (primitive endoderm marker, pre-implantation). β-catenin was detected in all cell lineages throughout pre-implantation development. Scale bar: 30 µm. **f**, Confocal immunofluorescence images of post-implantation embryos (E5.5 and E6.5) stained for β-catenin and SOX17 (visceral endoderm marker, post-implantation). β-catenin remained detectable in the post-implantation epiblast (Epi) and extraembryonic ectoderm (ExE). Scale bar: 30 µm. **g**, Confocal immunofluorescence images of blastocysts (∼E4.25) stained for Plakoglobin, OCT4 (epiblast marker) and SOX17 (Primitive endoderm marker). Fluorophore intensities were quantified along the white line marked in the merged image and are shown on the right. Primitive endoderm cells lost Plakoglobin expression once sorted to the inner cell mass’ surface. Scale bar: 30 µm. **h**, Confocal immunofluorescence images of human cleavage stage embryos stained for Plakoglobin and β-catenin. Scale bar: 50 µm. **i**, Confocal immunofluorescence images of a human blastocyst (Day 7) stained for SOX2 (Epiblast marker), SOX17 (Hypoblast marker) and Plakoglobin. Fluorophore intensities were quantified along the white line and are shown on the right. N=4 (remaining embryos shown in S4). Scale bar: 50 µm.

### Plakoglobin overexpression promotes the naïve pluripotency gene regulatory network

Considering that Plakoglobin was one of the most differentially expressed genes in microgel-cultured ESCs, we sought to test whether Plakoglobin overexpression could mimic the pluripotency-promoting effects of the microgel suspension culture system. We generated Plakoglobin-overexpressing cells by genomic integration of a *Jup-2A-mCherry* cassette under the control of a constitutive promoter in the RGd2 reporter line using the piggyBac™ system (**Figure 5A**). The resulting cell lines indicated exogeneous Plakoglobin expression via *Jup-2A-mCherry* and naïve pluripotency via *Rex1::GFPd2*. We derived clonal ESC lines overexpressing Plakoglobin at high (PG^HIGH^, clones #1-3) and low (PG^LOW^, clones #1-3) levels (**Figure 5B-C, S5A**). Consistent with previous reports^39^, we found a reciprocal correlation between Plakoglobin and β-catenin levels (**Figure 5D**). Furthermore, PG^HIGH^ cells showed higher nuclear Plakoglobin signal compared to PG^LOW^ cells (**Figure 5D, S5B**).

**Fig. 5.**
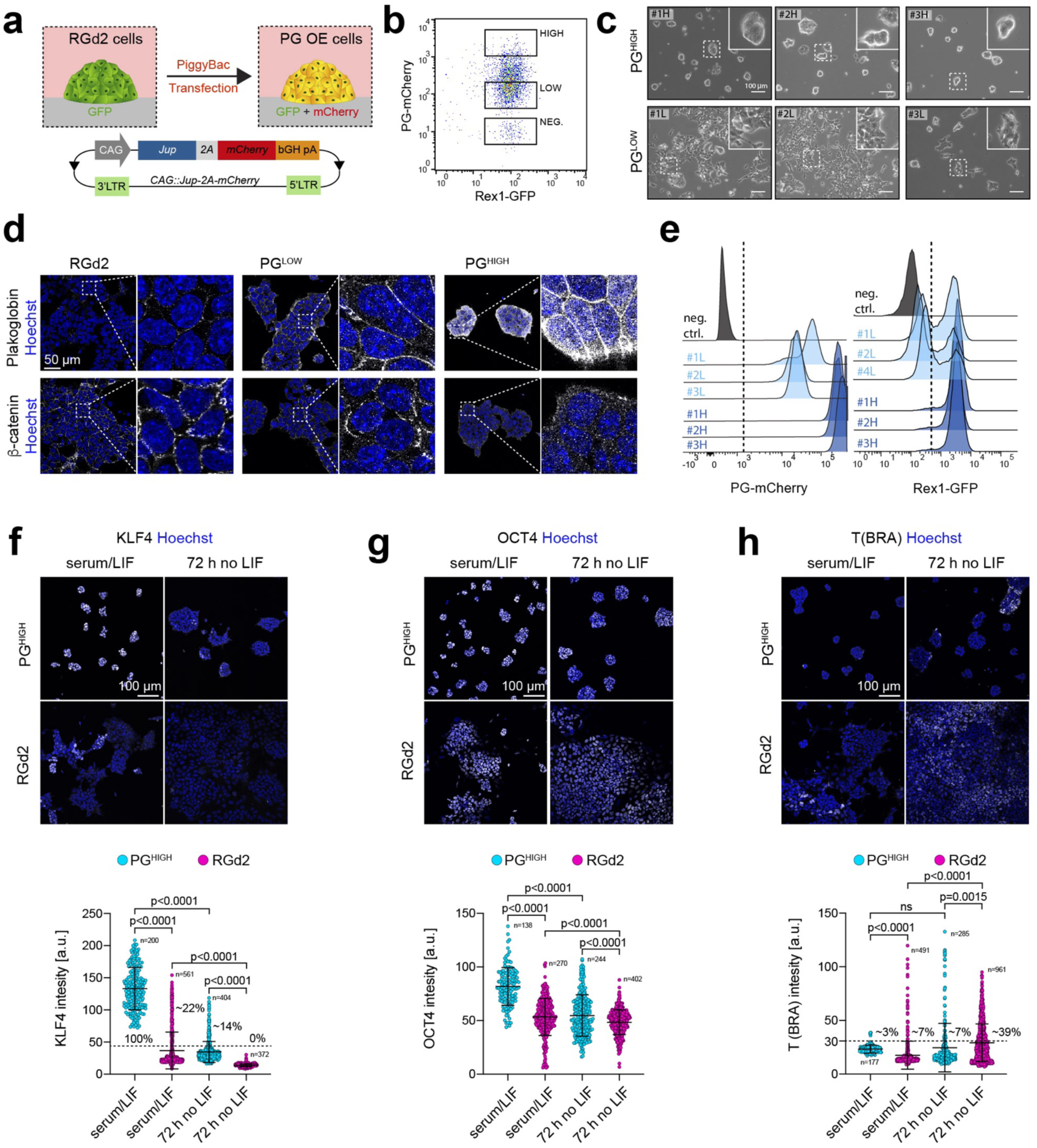
Plakoglobin expression promotes naïve pluripotency in meta-stable culture conditions. **a**, Schematic of the generation of Plakoglobin overexpressing RGd2 ESCs (PG OE). *Jup* (Plakoglobin) expression was placed under the control of a constitutively active CAG promoter and coupled to an mCherry fluorophore via a 2A self-cleaving peptide (*CAG::Jup-2A-mCherry*). This cell line simultaneously allowed monitoring of pluripotency via the Rex1-GFPd2 reporter and indicated the level of Plakoglobin via the mCherry signal. bGH pA: bovine growth hormone polyadenylation signal, LTR: long terminal repeats. **b**, Flow cytometric analysis of PG OE ESCs. Single cells were sorted to generate clonal populations of ESCs with high (PG^HIGH^) or low (PG^LOW^) levels of Plakoglobin overexpression. **c**, Phase contrast images of clonally expanded PG^LOW^ (#1L, #2L, #3L) and PG^HIGH^ (#1H, #2H, #3H) cells cultured in serum/LIF. Scale bar: 100 µm. **d**, Confocal immunofluorescence images of RGd2, PG^LOW^#2 and PG^HIGH^#3 cells stained for Plakoglobin (PG) and β-catenin. Scale bar: 50 µm. **e**, Flow cytometric analysis of the established clonal cell lines PG^LOW^ (#1L, #2L, #3L) and PG^HIGH^ (#1H, #2H, #3H) when cultured in serum/LIF. All PG^HIGH^ clones displayed a homogeneous Rex1-GFPd2 signal indicating acquisition of naïve-like pluripotent state. **f-h**, Confocal immunofluorescence images of RGd2 and PG^HIGH^ cells cultured in metastable pluripotent (serum/LIF) conditions and after 72 h of LIF withdrawal stained for the naïve pluripotency marker KLF4 (f), the general pluripotency marker OCT4 (h) and the gastrulation marker T(BRA) (h). Scale bar: 100 µm. Bottom: Nuclear fluorophore intensity quantification. For KLF4 and T (BRA), positive cells (KLF4 threshold: 45 a.u.; T (BRA) threshold: 30 a.u.) are indicated in [%].

We examined the ability of Plakoglobin-overexpressing ESCs to sustain features of naïve pluripotency by exposing the cells to metastable serum/LIF conditions. All PG^HIGH^ clones cultured in serum/LIF readily sustained the tightly packed, dome-shaped morphology associated with naïve pluripotency seen in 2i/LIF (**Figure 5C, S5A**). PG^LOW^ clones remained mostly as a monolayer (**Figure 5C**), similar to control RGd2 cells **(Figure S5E**). Strikingly, flow cytometric analysis revealed that all PG^HIGH^ clones displayed a uniformly positive Rex1-GFPd2 signal, indistinguishable from naïve ESCs cultured in 2i/LIF (**Figure 5E, S5A**). In contrast, PG^LOW^ cells maintained a bimodal GFP distribution, unless cultured in 2i/LIF (**Figure 5E, S5A**). We also assessed the naïve pluripotency markers KLF4 and TFCP2L1 (**Figure 5F, S5C**), the core pluripotency factor OCT4 (**Figure 5G**) and the gastrulation marker T (Brachyury) (**Figure 5H**) in the presence or absence of LIF. Consistent with the homogeneous Rex1-GFPd2 signal, we found that 100% of PG^HIGH^ cells in serum/LIF expressed the naïve pluripotency factor KLF4 in comparison to ∼22% of parental RGd2 cells (**Figure 5F**). Upon LIF removal for 72 h, these values dropped to ∼14% and 0%, respectively. The general pluripotency factor OCT4 was expressed in all conditions and slightly elevated in PG^HIGH^ cells (**Figure 5G**). The gastrulation marker T (Brachyury) was expressed in a subset (7% of cells) of serum/LIF cultured RGd2 cells and increased (39% of cells) after LIF removal (**Figure 5H**). In contrast, only 3% of PG^HIGH^ cells in serum/LIF expressed T (Brachyury), increasing to 7% after LIF removal. We repeated all experiments with polyclonal lines to exclude potential artifacts from clonal selection (**Figure S5D-M**). These results demonstrate that high Plakoglobin expression stabilizes the naïve pluripotency network in metastable culture conditions (serum/LIF), thereby recapitulating the effects of microgel suspension culture.

To comprehensively evaluate the effects of Plakoglobin overexpression on the pluripotency gene regulatory network, we performed single-cell RNA-seq using the inDrop workflow^40^ for the following samples: RGd2 ESCs in 2i/LIF, RGd2 ESCs in serum/LIF, encapsulated RGd2 ESCs in serum/LIF, clonal PG^LOW^ ESCs in serum/LIF and clonal PG^HIGH^ ESCs in serum/LIF (**Figure 6A**). Pearson correlation coefficient computation between all samples revealed that PG^HIGH^ ESCs in serum/LIF were transcriptionally most similar to the naïve control cells in 2i/LIF (**Figure 6B**). All remaining serum/LIF samples exhibited lower degrees of resemblance with naïve control cells (RGd2 ESCs in 2i/LIF). Principal component analysis separated pluripotent cell populations (RGd2 ESCs in 2i/LIF, PG^HIGH^ ESCs in serum/LIF, encapsulated RGd2 ESCs in serum/LIF) from differentiating cells (PG^LOW^ ESCs in serum/LIF, RGd2 ESCs in serum/LIF) along the first principal component (**Figure 6C**). We determined developmental trajectories by UMAP (**Figure 6D**) and RNA latent time computation using dynamical RNA velocity as an input^41, 42^ (**Figure 6D-I**), which revealed the presence of a continuum of cell states from naïve to primed pluripotency (**Figure 6D-I**). The core pluripotency factor *Pou5f1* (OCT4) was expressed in most cells, whereas expression of naïve transcription factors including *Esrrb*, *Klf2*, *Tfcp2l1* and *Zfp42* was restricted to PG^HIGH^ cells and the 2i/LIF control (**Figure 6E-F**). The early post-implantation markers *Pou3f1* (OCT6) and *Otx2*^10, 43^ and early differentiation markers such as *Anxa2*, *Krt18* and *Tubb6*^44^ were upregulated in RGd2 ESCs on plastic and PG^LOW^ ESCs in serum/LIF conditions, but absent from RGd2 ESCs in agarose microgels (**Figure 6G**). PG^HIGH^ differed from all other serum/LIF cultured samples by its upregulation of naïve pluripotency-associated threonine dehydrogenase (*Tdh*)^45, 46^ and downregulation of γ-actin (*Actg1*) (**Figure S6B-D**). 2i/LIF control cells separated from all serum/LIF samples (including PG^HIGH^ cells) by their upregulation of *Vim*, *Scd2*, *Ldhb* and downregulation of *Tmsb4x*, *Mycn* and *Ldha* (**Figure S6E-F)**. These data demonstrate that high Plakoglobin expression is more effective in sustaining naïve pluripotency compared to microgel encapsulation alone, which mainly prevents the upregulation of early lineage specifiers.

**Fig. 6.**
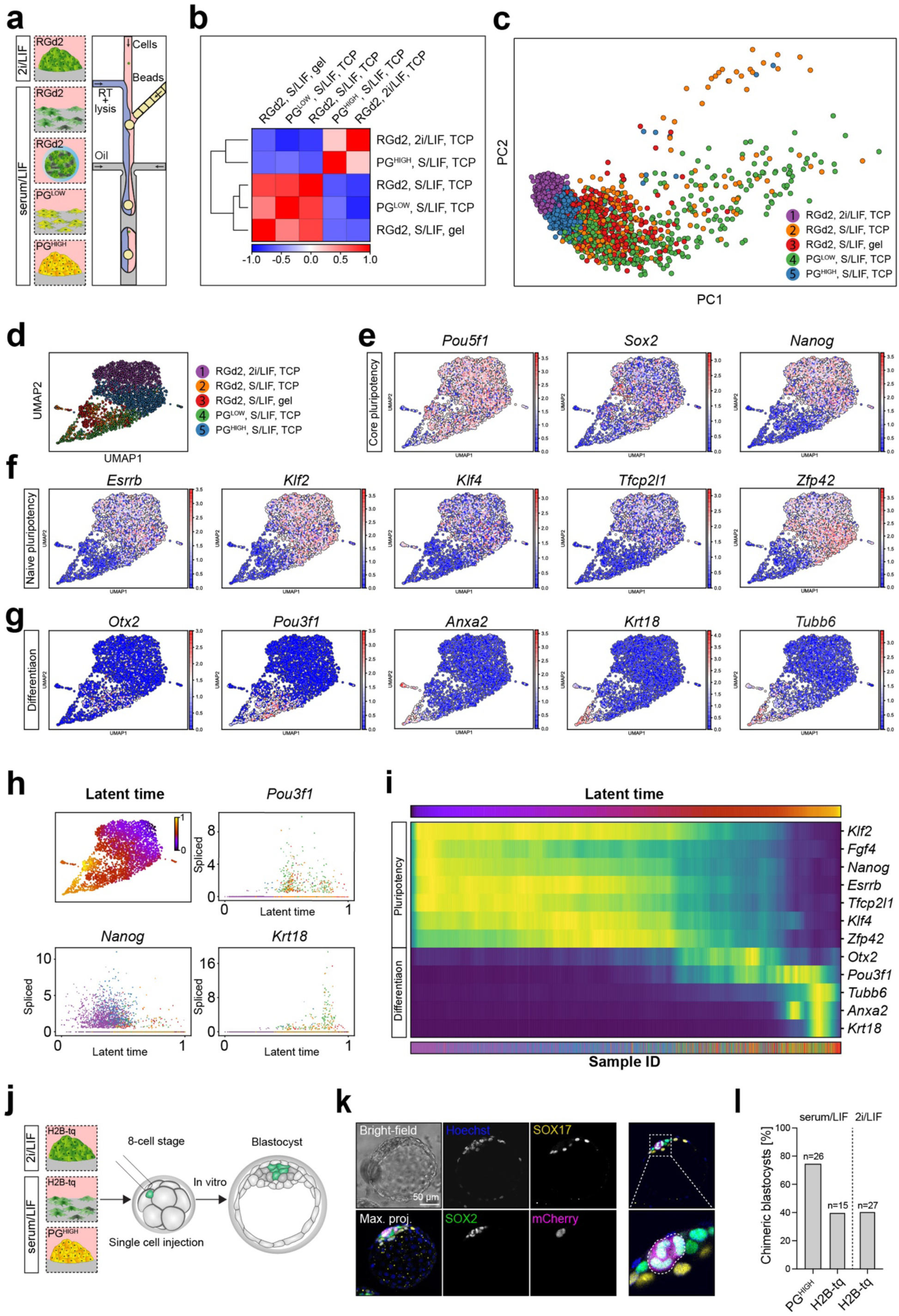
Single-cell sequencing elucidates plakoglobin-induced re-establishment of naïve pluripotency. **a**, Schematic of the different samples subjected to scRNA-seq analysis using the inDrop workflow. (RT: reverse transcriptase). **b**, Pearson correlation heatmap highlighting the resemblance between the PG^HIGH^ ESCs cultured under metastable pluripotent conditions (serum/LIF) and the control RGd2 ESCs cultured under naïve pluripotent (2i/LIF) conditions. Samples with low (PG^LOW^) or no Plakoglobin over-expression as well as cells encapsulated in microgels in serum/LIF were highly inter-correlated but bore little correspondence with the two other conditions. **c**, Principal component (PC) analysis illustrating the spread of the single-cells across PC1. The naïve pluripotent (2i/LIF) control RGd2 cells and the PG^HIGH^ (in serum/LIF) samples clustered together and showed little spread across the PC1 axis indicating similar and homogeneous gene expression, respectively. By contrast to PG^HIGH^ cells, the remainder of serum/LIF cultured cells (PG^LOW^, RGd2 on plastic and RGd2 in microgel) clustered apart from the naïve control cells and spread heavily across the PC1 axis, underlining higher levels of heterogeneity in gene expression. **d**, Uniform Manifold Approximation and Projection (UMAP) dimensional reduction plot of the scRNA-seq samples, for the maps of individual markers (in E, F and G). **e**, Gene-expression values projected on the UMAP plot for core pluripotency markers (*Pou5f1*, *Sox2* and *Nanog*). **f**, Gene-expression values projected on the UMAP plot for naïve pluripotency markers (*Esrrb*, *Klf2*, *Klf4*, *Tfcp2l1* and *Zfp42*), highlighting an up-regulation in the PG^HIGH^ and naïve pluripotent (2i/LIF) conditions. **g**, Gene-expression values projected on the UMAP plot for peri-implantation (*Otx2* and *Pou3f1*) and serum-induced differentiation markers (*Anxa2*, *Krt18* and *Tubb6*), highlighting an up-regulation in a fraction of the serum/LIF samples with the exception of the PG^HIGH^ clone which did not harbor hallmarks for differentiation. **h**, (top left) Latent time computation generated using scVelo and projected on the UMAP underlining the continuum between naïve pluripotency (latent time = 0) and differentiation (latent time = 1). (top right) Single-cell spliced read counts arranged across the computed latent time for *Pou3f1* (top right), *Nanog* (bottom left) and *Krt18* (bottom right). **i**, Heatmap representing the changes in gene expression across all cells arranged by latent time progression, further depicting the transition from naïve pluripotency to differentiation. **j**, Schematic of single-cell injection into 8-cell stage embryos with subsequent *in vitro* culture until the blastocyst stage. **k**, Confocal immunofluorescence image of a chimeric blastocyst that was injected at the 8-cell stage with a single serum/LIF cultured PG^HIGH^ ESC. Blastocysts were stained for SOX2 (epiblast marker) and SOX17 (primitive endoderm marker). PG^HIGH^ cells were identified by the PG-mCherry signal as shown in the merged image and highlighted by the white dotted line. Scale bar: 50 µm. **l**, Chimeric blastocyst contribution efficiency of PG^HIGH^ cells in serum/LIF and control H2B-tq cells in naïve (2i/LIF) and metastable pluripotent (serum/LIF) conditions. Single cells were injected at the 8-cell stage followed by *in vitro* culture until blastocyst stage.

To functionally assess the self-renewal and developmental potential of serum/LIF cultured PG^HIGH^ ESCs, we compared RGd2 ESCs against PG^HIGH^ ESCs using an *in vitro* clonogenicity and single-cell chimeric contribution assays. PG^HIGH^ cells displayed high colony forming efficiencies in 2i/LIF and serum/LIF, while the self-renewal capacity of serum/LIF cultured RGd2 ESCs was reduced upon replating into naïve 2i/LIF culture conditions (**Figure S6A**). To examine the ability of PG^HIGH^ ESCs to contribute to the blastocyst, we injected individual PG^HIGH^ ESCs cultured in serum/LIF, control H2B-tq (serum/LIF) and naïve H2B-tq (2i/LIF) ESCs into 8-cell stage embryos and cultured them *in vitro* until the blastocyst stage (**Figure 6J**). The single cell injections resulted exclusively in contribution to the epiblast, but not extraembryonic lineages (**Figure 6K**). Remarkably, PG^HIGH^ cells exhibited ∼1.8-fold increased chimeric blastocyst contribution efficiency (∼77%) compared to the wildtype cells in 2i/LIF (∼41%) and serum/LIF (∼40%) (**Figure 6L**). Taken together, with high levels of exogenous Plakoglobin expression ESCs acquire a transcriptional state reminiscent of naïve pluripotency even under serum/LIF conditions and cells in this state readily contribute to the developing blastocyst.

### Plakoglobin maintains naïve pluripotency independently of β-catenin

To determine how Plakoglobin sustains naïve pluripotency, we interrogated the signaling requirements of Plakoglobin-overexpressing ESCs for self-renewal (**Figure 7A**). In 2i/LIF, the combination of any two of the three components (PD, CH and LIF) is sufficient to maintain naïve pluripotency^20^. We challenged self-renewal for more than five passages of wildtype (RGd2), PG^LOW^ and PG^HIGH^ ESCs in medium supplemented with only one of the individual 2i/LIF components (**Figure 7B-D**). Naïve pluripotency was assessed through the Rex1-GFPd2 reporter by flow cytometry. In the presence of LIF-only, wildtype (RGd2) ESCs could not be passaged beyond day 6, consistent with previous observations^16^. PG^LOW^ displayed reduced Rex1-GFPd2 levels, but could be propagated (**Figure 7B, F**). Remarkably, PG^HIGH^ ESCs cultured with LIF-only gave rise to dome-shaped morphology and displayed homogenous (>99%) Rex1-GFPd2 levels, indistinguishable from 2i/LIF cells (**Figure 7B, E**). In PD-only conditions, wildtype (RGd2) and PG^LOW^ ESCs were unable to self-renew and vanished by day 4 (**Figure 7C, F**). PG^HIGH^ ESCs exhibited robust expression of Rex1-GFPd2 (∼75-90%) and could be propagated throughout the time course, suggesting that the rescue effect of Plakoglobin expression was further amplified in the presence of PD (**Figure 7C, E**). Wildtype (RGd2) cultured in the presence of CH-only exhibited medium (>60%) Rex1-GFPd2 levels at day 6 and could not be stably propagated (**Figure 7D**). PG^LOW^ ESCs showed some heterogeneity, but could be passaged until day 16 (**Figure 7D, F**). Notably, PG^HIGH^ ESCs failed to maintain pluripotency (**Figure 7F**). Collectively, the single-component time course analysis demonstrated that Plakoglobin overexpression is capable of sustaining naïve pluripotency in combination with PD or LIF, but not CH. To find out whether Plakoglobin can functionally replace β-catenin to maintain naïve pluripotency, we cultured ESCs in the presence of the tankyrase inhibitor XAV939 (XAV) to abrogate nuclear β-catenin signaling^47^. We performed single-component (PD, CH or LIF) time course analysis of PG^HIGH^ and PG^LOW^ ESCs in the presence of XAV. Self-renewal with LIF-only or PD-only in PG^HIGH^ cells was unaffected by XAV treatment. Only PG^LOW^ cells supplemented with CH, started to differentiate and lost GFP expression upon XAV treatment (**Figure 7G**). To confirm that Plakoglobin-mediated maintenance of pluripotency is truly independent of β-catenin, we generated two clonal *Ctnnb1* (β-catenin) knockout PG^HIGH^ cell lines: PG^HIGH^#3 *Ctnnb1* KO#2 and PG^HIGH^#3 *Ctnnb1* KO#12, both of which remained naïve pluripotent in 2i/LIF (**Figure 7H, S7A-D**). Remarkably, PG^HIGH^ *Ctnnb1*-KO cells supplemented with either LIF-only or PD-only remained naïve pluripotent with near homogeneous Rex1-GFPd2 expression. In contrast, PG^HIGH^ *Ctnnb1*-KO supplemented with CH-only failed to maintain pluripotency and died (**Figure 7I, 7J**). These data establish that the naïve pluripotent circuitry is supported by Plakoglobin, independently of β-catenin.

**Fig. 7.**
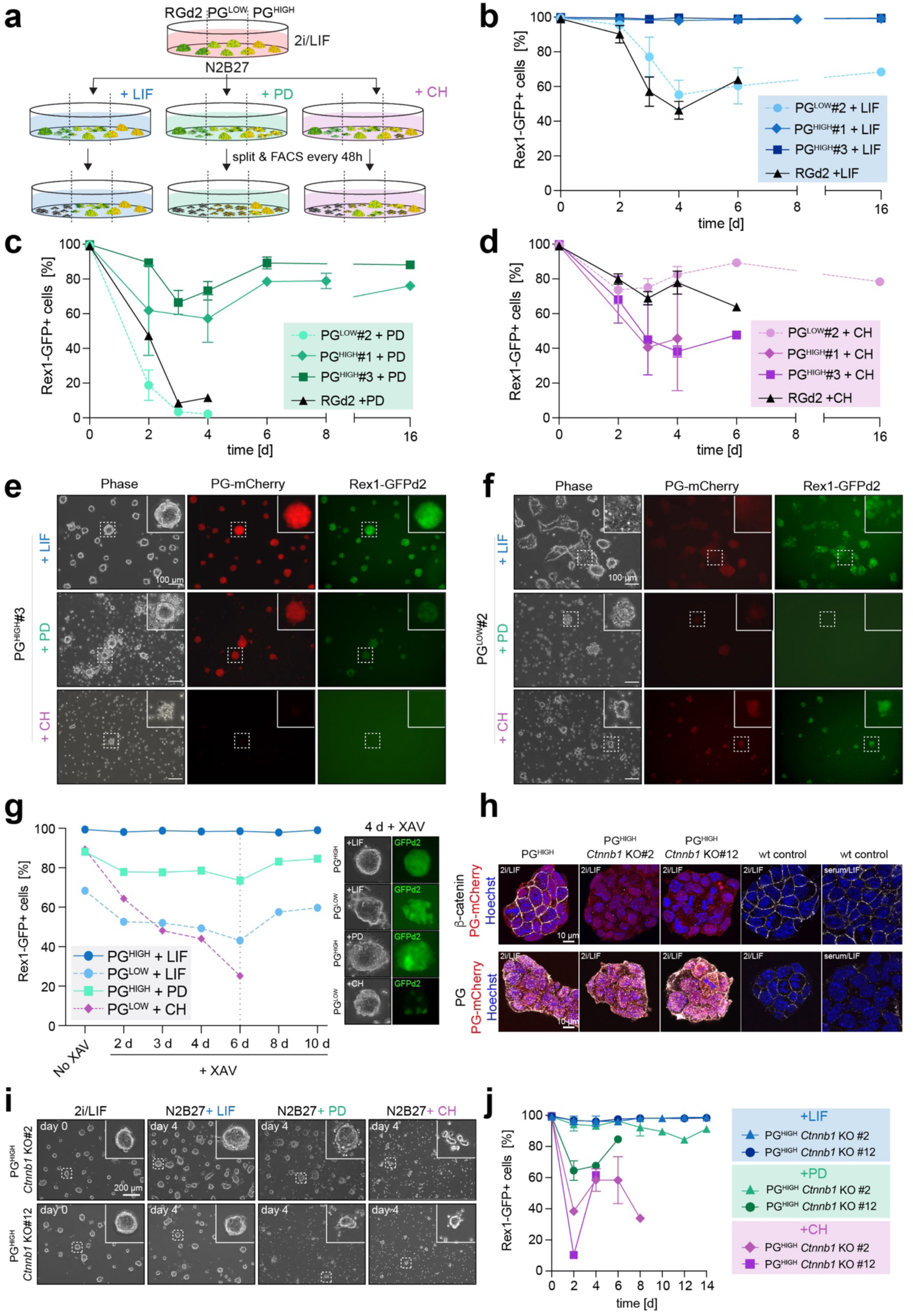
Plakoglobin sustains pluripotency independently of β-catenin. **a**, Experimental layout of culture conditions to asses Plakoglobin’s potential to maintain naïve pluripotency. PD (MEK inhibitor) CH (WNT agonist). **b-d**, Flow cytometric analysis of the Rex1-GFPd2 reporter in the PG^HIGH^ (clone #1 and #3), PG^LOW^ and RGd2 cell lines. Cells were cultured up to 16 days with the sole supplementation of PD (b), LIF (c) or CH (d). **e-f**, Phase contrast and epifluorescence (PG-mCherry and Rex1-GFPd2) images of PG^HIGH^ and PG^LOW^ cells 6 days after culture in either PD, LIF or CH. Scale bar: 100 µm. **g**, Flow cytometric analysis of the Rex1-GFPd2 reporter in the surviving cell lines (established from B-D) after treatment with the WNT-signaling antagonist XAV. Cells were measured every 48 hours for 10 days. (Right) Phase contrast and epifluorescence (Rex1-GFPd2) representative images of the cells after 4 days of XAV treatment. **h**, Confocal immunofluorescence images of PG^HIGH^, PG^HIGH^ *Ctnnb1* KO (clone #2 and #12) and wild type control cells stained for β-catenin and Plakoglobin. Scale bar: 10 µm. **i**, Phase contrast images of PG^HIGH^ *Ctnnb1* KO (clone #2 and #12) cells cultured in 2i/LIF or with the single supplementation of LIF, PD or CH for 4 days. Scale bar: 200 µm. **j**, Flow cytometric analysis of the Rex1-GFPd2 reporter in PG^HIGH^ *Ctnnb1* KO (clone #2 and #12) cells when cultured with the sole supplementation of LIF, PD or CH. Cells were analyzed every 48 hours for up to 14 days.

## Discussion

It has become increasingly clear that cell behavior and developmental processes are not just regulated via biochemical processes but also succumb to a plethora of mechanical signals, however the interdependence of these levels of control remains underdetermined^48, 49^. Conventional 2D plastic dishes do not emulate the complex *in vivo* environment and are unsuitable to test for and decipher cell signaling in 3D. New experimental formats are needed to shed light on how gene regulatory networks, biochemical and biomechanical signals are integrated within cells. Here we used a microgel system to encapsulate embryonic stem cells into spherical 3D agarose scaffolds. Consistent with previous studies, we found the soft, non-degradable 3D matrix to be beneficial for pluripotency^21, 25, 50, 51^. The interpretation of RNA-seq data on microgel-encapsulated cells revealed Plakoglobin, a vertebrate homologue of β-catenin and a crucial factor for embryonic development, as one of the most differentially upregulated genes^52, 53^. Both proteins are known to be involved in embryonic development: while β-catenin is a well-known mechanotransducer in several species and cell types^4, 54–56^, Plakoglobin has so far only been shown to be recruited to adherens junctions under tension in *Xenopus* mesendoderm cells^57^. To our knowledge, little is known about Plakoglobin’s ability to react to mechanical signals in mammals – in particular during pluripotency. Remarkably we observed that Plakoglobin was exclusively upregulated in microgels but not in hanging drops (3D environment without confinement) or on soft 2D microgels. We therefore suggest that microgel-mediated confinement acts as an upstream mechanical signal for Plakoglobin expression. Notably, Plakoglobin’s expression *in vivo* correlates with the expansion of the blastocyst’s cavity and thus the increase in pressure within the blastocyst, an expression pattern we observed in mouse and human embryos^31, 32, 58^. Yet, whether there is a mechanoresponsive component of Plakoglobin regulation *in vivo* remains to be investigated. Simultaneously with the expression of Plakoglobin in the pre-implantation epiblast, naïve pluripotency is established^9, 58^. This observation and the abrupt disappearance of Plakoglobin upon implantation (and transition into primed pluripotency) suggest a role for Plakoglobin in specifically regulating naïve pluripotency. Plakoglobin, unlike β-catenin, is known to activate TCF/LEF-mediated transcription only weakly^59, 60^ and endogenous levels of plakoglobin cannot rescue β-catenin null embryos during gastrulation^61, 62^. Yet, Plakoglobin has been shown to delay differentiation, but its ability to maintain and regulate pluripotency is not known^59^. Here, comprehensive single-cell sequencing and blastocyst injections confirmed that Plakoglobin overexpressing cells re-acquire naïve-like pluripotency under the metastable pluripotent culture conditions of serum/LIF. In *Xenopus* it has been suggested that Plakoglobin could act indirectly through β-catenin by releasing it from the membrane and saturating its degradation machinery whilst another study found that in mammalian cells plakoglobin expression lead to the degradation of β-catenin^63^. Here, XAV treatment (blocking nuclear translocation of β-catenin) or a β-catenin knockout in Plakoglobin-overexpressing cells does not compromise their ability to maintain the naïve pluripotency network. This suggests, that Plakoglobin can act independently of β-catenin in supporting pluripotency. Whether Plakoglobin’s beneficial effect on pluripotency acts in a redundant fashion to β-catenin, by alleviating repressive effects of TCF7L1 on the pluripotency network, remains to be investigated^16, 17, 64, 65^. In summary, the agarose microgel format allowed us to investigate the effects of microenvironmental confinement on pluripotent ESCs. We identified Plakoglobin (*Jup*) expression to be confinement-induced, a mechanism that would have been missed under conventional cell culture conditions. Furthermore, owing to Plakoglobin’s expression in human and mouse pre-implantation embryos – in particular the expression in the epiblast, and its ability to reinstate naïve pluripotency in ESCs – led us to propose Plakoglobin as an evolutionarily conserved, mechanoresponsive regulator of naïve pluripotency. More generally, our approach of using agarose microgels may be instructive to elucidate confinement-mediated signaling cascades beyond pluripotent stem cells, e.g. in developmental processes such as gastrulation and diseases including cancer – as all of these have been postulated to involve mechanosensitive signaling pathways^66, 67^.

## Acknowledgements

T.N.K. and M.H. received scholarship support from AstraZeneca, J.D.J. from the BBSRC, A.L.E. from the Cambridge Trusts and the EU H2020 Marie Curie ITN MMBio and K.F. from the MRC and St. John’s College Cambridge. This work was supported by the Wellcome Trust (WT108438/C/15/Z). F.H. is an H2020 ERC Advanced Investigator (695669). We are grateful to Dr. Geraldine M. Jowett for critical reading and constructive comments on the manuscript.

## Author contribution

T.N.K. and F.H. conceived and initiated the study. T.N.K. and A.L.E. carried out experiments across the entire range of approaches, analyzed data, prepared figures, and organized the preparation of the manuscript. J.D.J. prepared the inDrop and bulk sequencing libraries and analyzed transcriptomic data. A.Y. performed the mouse embryo injections. J.N. holds the required HFEA and Home Office licenses and carried out the human embryo experiments. M.H. and K.F. assisted with generation of cell lines. A.W. and Kr.F. implemented AFM measurements and their analysis. E.M.S., C.R. and S.B. contributed exploratory data and C.M. and K.C. experimental approaches. T.N.K., T.E.B. and F.H. wrote the manuscript with input from all authors. T.E.B. and F.H. supervised the work.

---

**Extended Data Fig. 1.**
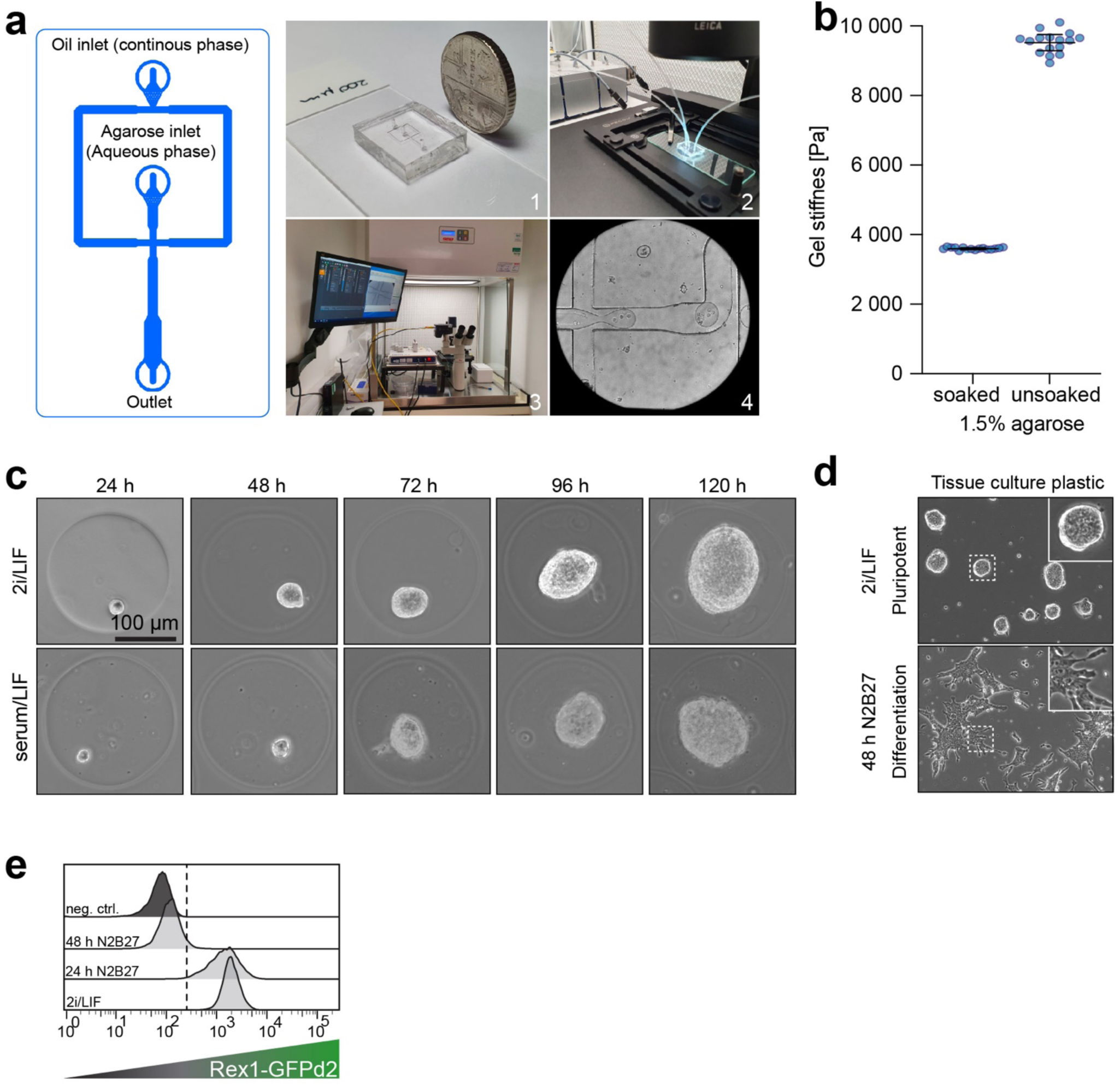
Microfluidic-based cell encapsulation. **a**, Schematic of the microfluidic flow-focusing device (FFD) and explanatory images of the encapsulation rig set-up: (1) Single-inlet FFD, (2) Fully connected FFD with oil and agarose inlet and outlet, (3) Microfluidic encapsulation rig in laminar flow hood and (4) FFD crossing of aqueous (agarose) and continuous (oil) phase. **b**, Measurements of the elastic modulus of the ultra-low melting agarose by atomic force microscopy. Measurements were performed on bulk gels that were pre-incubated with medium (soaked) or without pre-incubation. **c**, Representative time course phase contrast images of encapsulated ESCs after 24 h, 48 h, 72 h, 96 h and 120 h of culture in naïve pluripotent (2i/LIF) and metastable pluripotent (serum/LIF) conditions. **d**, Phase contrast images of ESCs cultured on conventional tissue culture plastic in 2i/LIF (pluripotent) and 48 h of culture in N2B27 (differentiation). **e**, Flow cytometric analysis of the RGd2 cells during pluripotency (2i/LIF) 24 h and 48 h of differentiation (N2B27). Cells have fully exited after 48 h of differentiation as indicated by the complete loss of Rex1-GFPd2.

**Extended Data Fig. 2.**
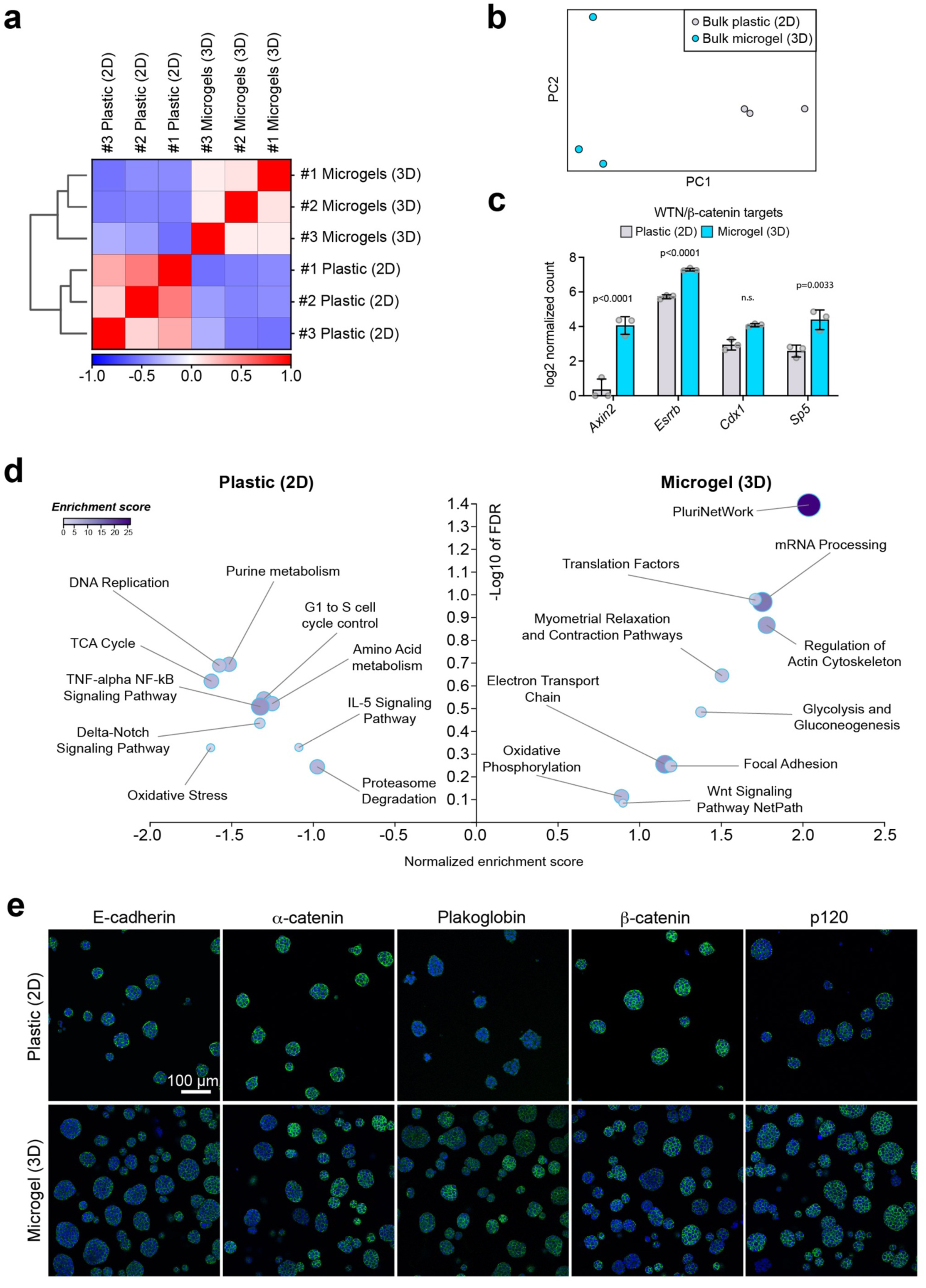
Comparison of cell culture in microgels (3D) and on plastic (2D). **a**, Pearson correlation coefficient heatmaps for each bulk RNA-seq replicate sample for both ESCs seeded on plastic (2D) or encapsulated in microgels (3D) under naïve pluripotent (2i/LIF) culture conditions. **b**, Principal component (PC) dimensional reduction plot of each sequenced replicate from ESCs cultured in 2i/LIF on plastic or encapsulated in microgels. **c**, Bulk RNA-seq analysis of known WNT/β-catenin target genes. Error bars indicate the mean and standard deviations. **d**, Gene set enrichment analysis using the most differentially expressed genes (|log2 fold-change| > 0.5 and Bonferroni-adjusted p-value < 10^-3) between ESCs cultured on tissue culture plastic in 2i/LIF and in agarose microgels, highlighting a significant enrichment of the PluriNetWork (WikiPathways) under microgel culture conditions. **e**, Confocal immunofluorescence images of naïve pluripotent (2i/LIF) cultured ESCs on plastic (2D) and encapsulated in microgels (3D). Cells were stained for E-cadherin, α-catenin, β-catenin, Plakoglobin and δ-catenin. Scale bar: 100 µm

**Extended Data Fig. 3.**
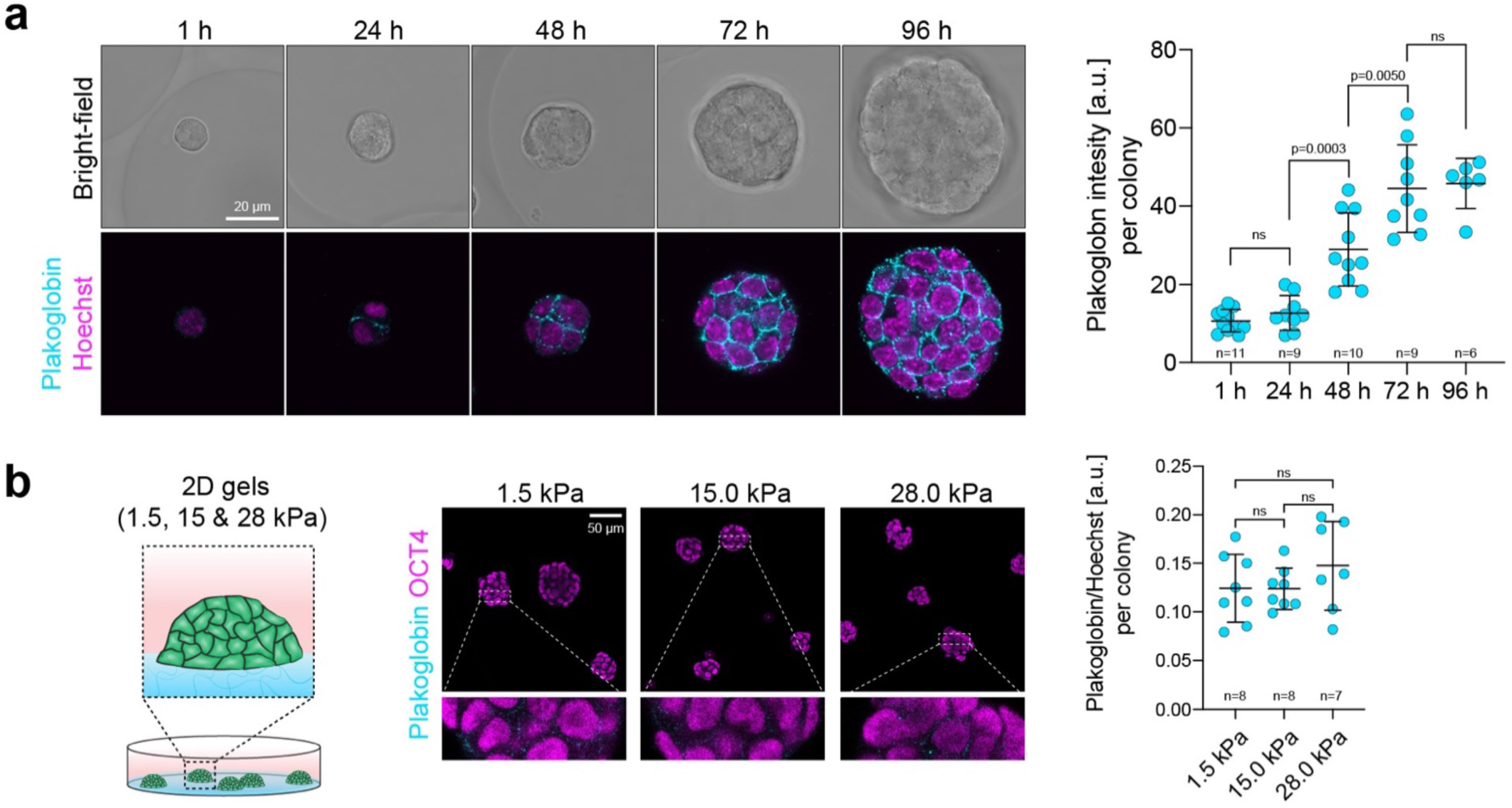
Microenvironmental regulation of Plakoglobin. **a**, Representative confocal immunofluorescence images of a time-course over 96 h of naïve pluripotent (2i/LIF) microgel encapsulated ESCs. Cells were stained for Plakoglobin. Plakoglobin levels were quantified per colony, significant upregulation was observed as early as 48 h after encapsulation. Error bars indicate the mean and standard deviations. **b**, Confocal immunofluorescence images of naïve pluripotent (2i/LIF) ESCs cultured on commercially available (see methods) 2D Polydimethylsiloxane gels with varying stiffnesses (1.5 kPa, 15 kPa and 28 kPa). Cells were stained for OCT4, F-actin and Plakoglobin. Plakoglobin levels (indicated on the right) remained constant irrespectively of 2D gel stiffness. Error bars indicate the mean and standard deviations.

**Extended Data Fig. 4.**
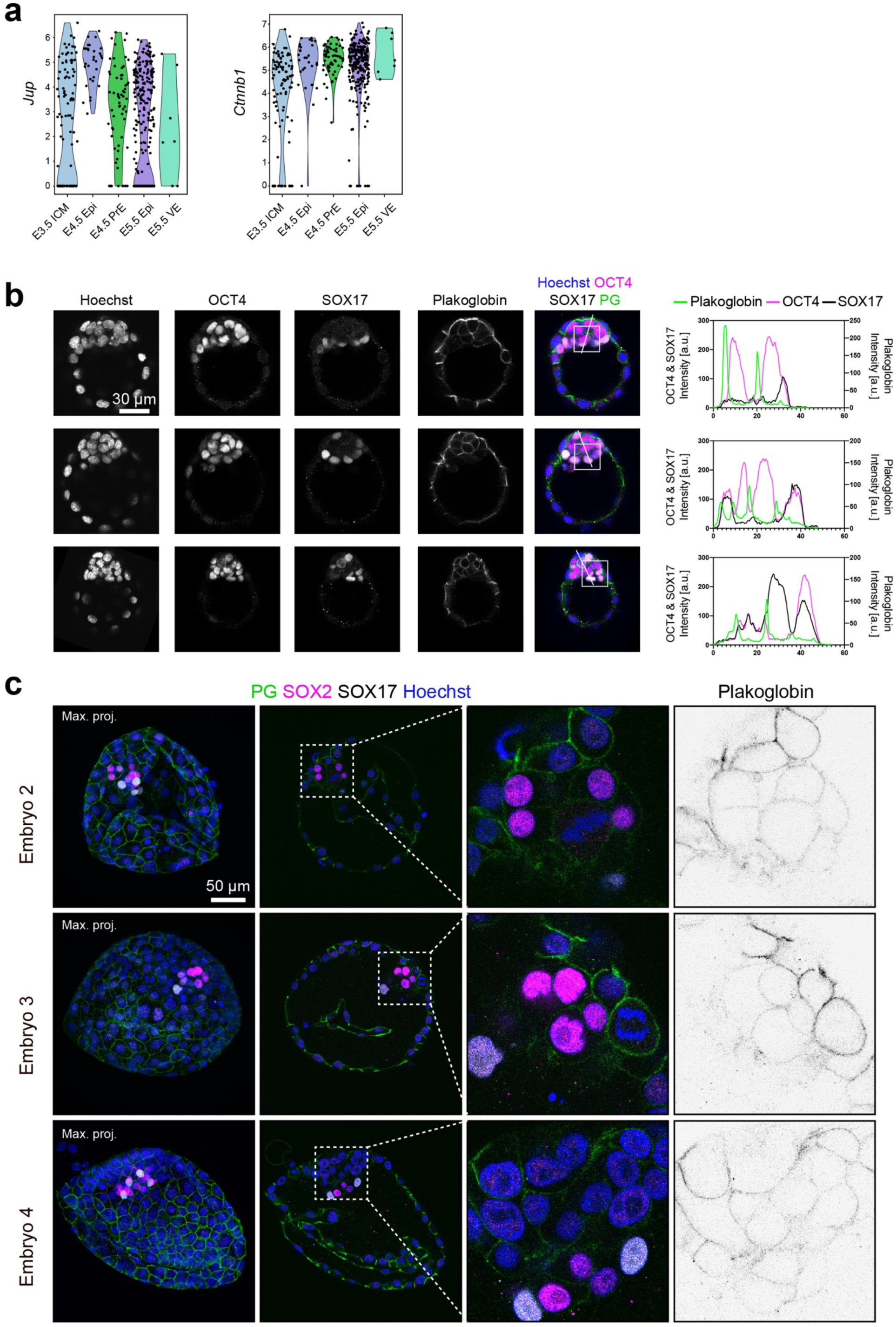
Plakoglobin expression in the embryo. **a**, Meta-analysis of single cell RNA-seq data^68^ from mouse embryos for *Jup* (Plakoglobin) and *Ctnnb1* (β-catenin) in log2 normalized counts during early embryogenesis. ICM = Inner Cell Mass, EPI = Epiblast, PrE = Primitive Endoderm, VE = Visceral Endoderm. **b**, Confocal immunofluorescence images of blastocysts (∼E4.25) stained for Plakoglobin, OCT4 (epiblast marker) and SOX17 (Primitive endoderm marker). Fluorophore intensities were quantified along the white line marked in the merged image and are shown on the right. Primitive endoderm cells lost Plakoglobin expression once sorted to the inner cell mass’ surface. Scale bar: 30 µm. **c**, Confocal immunofluorescence images of human blastocysts (Day 7) stained for SOX2 (Epiblast marker), SOX17 (Hypoblast marker) and Plakoglobin. Fluorophore intensities were quantified along the white line and are shown on the right. Scale bar: 50 µm.

**Extended Data Fig. 5.**
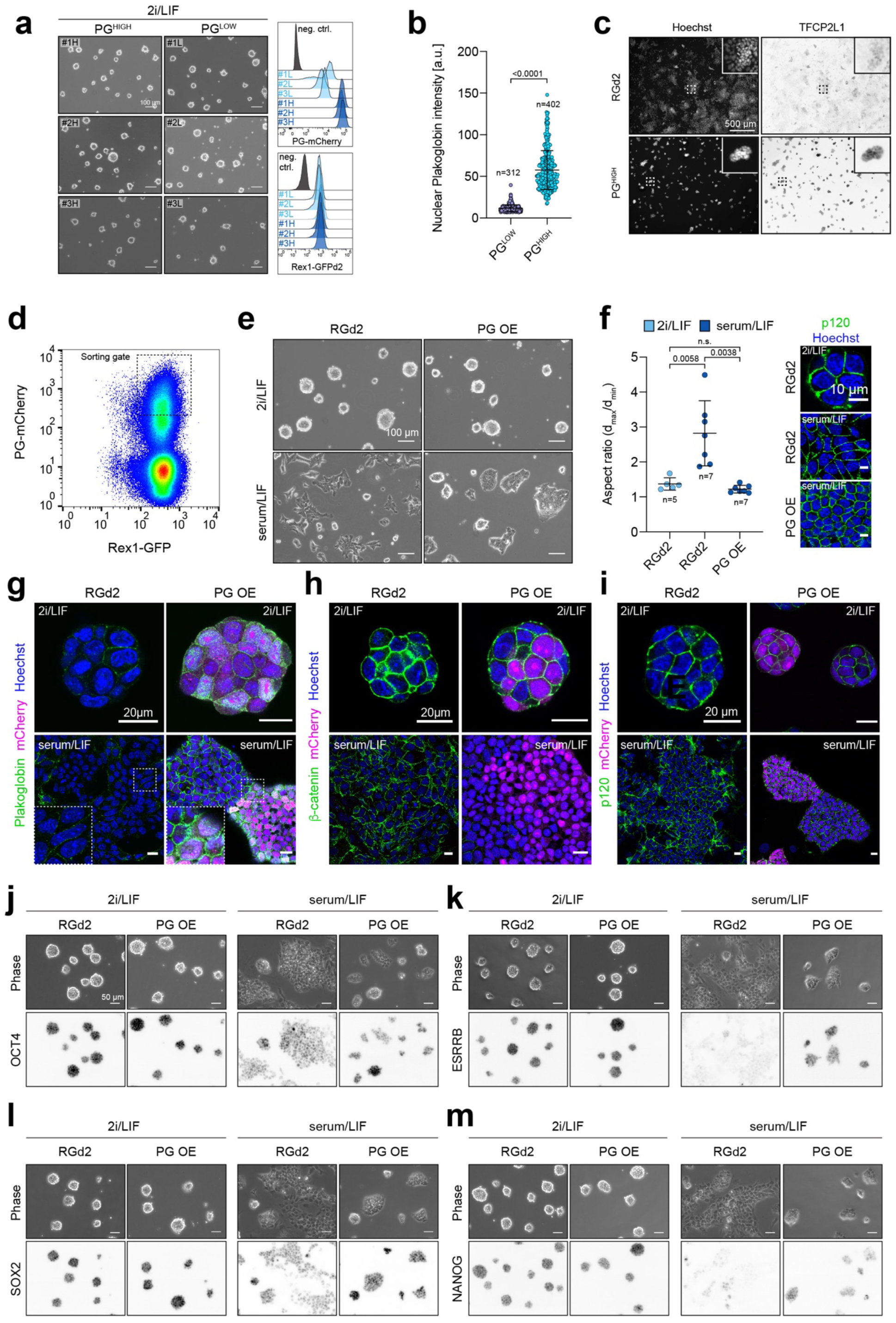
Plakoglobin expression promotes naïve pluripotency in meta-stable culture conditions. **a**, Phase contrast images of clonally expanded PG^LOW^ (#1L, #2L, #3L) and PG^HIGH^ (#1H, #2H, #3H) cells cultured in 2i/LIF. Scale bar: 100 µm. **b**, Quantification of nuclear Plakoglobin immunofluorescence signal (ref. Fig.5D) in PG^LOW^ (n = 312) and PG^HIGH^ (n = 402) cells cultured under serum/LIF conditions. Error bars indicate the mean and standard deviations. **c**, Epifluorescence images of RGd2 and PG^HIGH^ cells in metastable pluripotent conditions (serum/LIF) stained for the naïve pluripotency marker TFCP2L1. Scale bar: 500 µm. **d**, Flow cytometric analysis of *CAG::Jup-2A-mCherry* transfected RGd2 cells. PO OE cells were sorted based on positive Rex1-GFPd2 and PG-mCherry signal (sorting gate indicated as a rectangle). **e**, Phase contrast images of RGd2 and PG OE cells cultured in naïve pluripotent (2i/LIF) and metastable pluripotent (serum/LIF) conditions. PG OE cells remained as tightly packed colonies under metastable conditions in serum/LIF similar to naïve cells in 2i/LIF. Scale bar: 100 µm. **f**, Aspect ratio measurements (distance_max_/distance_min_) of RGd2 cells (in 2i/LIF n = 5, in serum/LIF n = 7) and PG OE cells (in serum/LIF n = 7). Cell morphology is indicated by confocal immunofluorescence images on the right with cells stained for membrane adjacent p120. Scale bar: 10 µm. **g-i**, Confocal immunofluorescence images of RGd2 and PG OE cells in naïve pluripotent (2i/LIF) and metastable pluripotent (serum/LIF) conditions. Cells were stained for Plakoglobin (d), β-catenin (e) and p120 (f). Scale bar: 20 µm. **j-m**, Epifluorescence and phase contrast images of RGd2 and PG OE cells under naïve pluripotent (2i/LIF) and metastable pluripotent (serum/LIF) conditions. Cells were stained for OCT4 (g), ESRRB (h), SOX2 (i) and NANOG (j). Scale bar: 50 µm.

**Extended Data Fig. 6.**
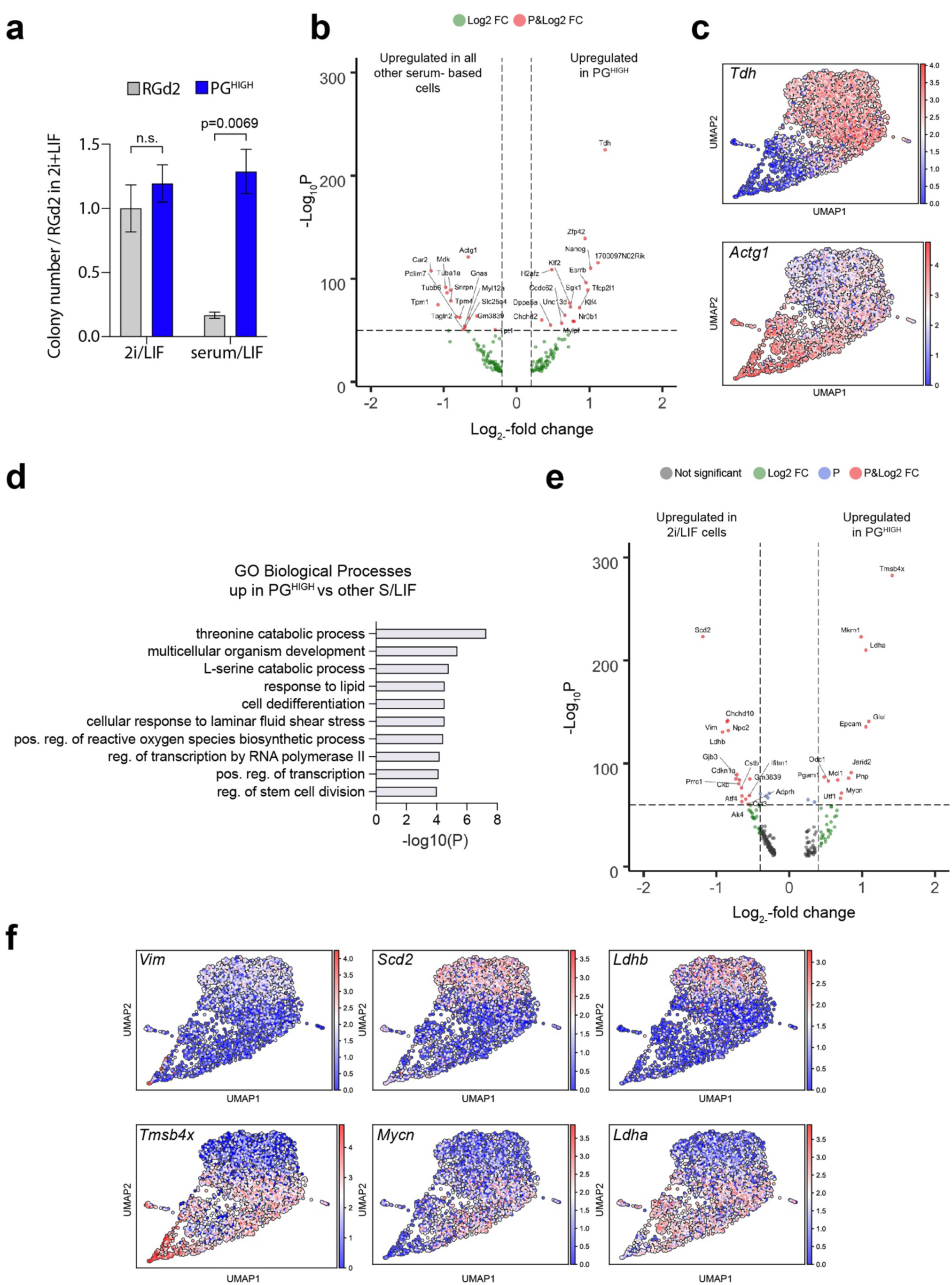
Single-cell sequencing elucidates plakoglobin-induced re-establishment of naïve pluripotency. **a**, Colony formation assay of RGd2 and PG^HIGH^ cells. (N = 3)**. b**, Volcano plot of genes with log2-fold change > 0.2 and -log10P > 50, comparing PG^HIGH^ cells against all other serum/LIF cultured samples. **c**, Gene-expression values projected on the UMAP plot for *Tdh* (upregulated in naïve 2i/LIF control cells and PG^HIGH^ in serum/LIF) and *Actg1* (upregulated in all other serum/LIF samples). **d**, List of Gene Ontology for Biological Processes upregulated in PG^HIGH^ cells compared to all other serum/LIF samples. **e**, Volcano plot of genes with log2-fold change > 0.4 and -log10P > 60, comparing PG^HIGH^ cells against naïve (2i/LIF) control cells. **f**, Gene-expression values projected on the UMAP plot for *Vim*, *Scd2* and *Ldhb* (upregulated in naïve 2i/LIF control cells) and *Tmsb4x*, *Mycn* and *Ldha* (upregulated in all serum/LIF samples).

**Extended Data Fig. 7.**
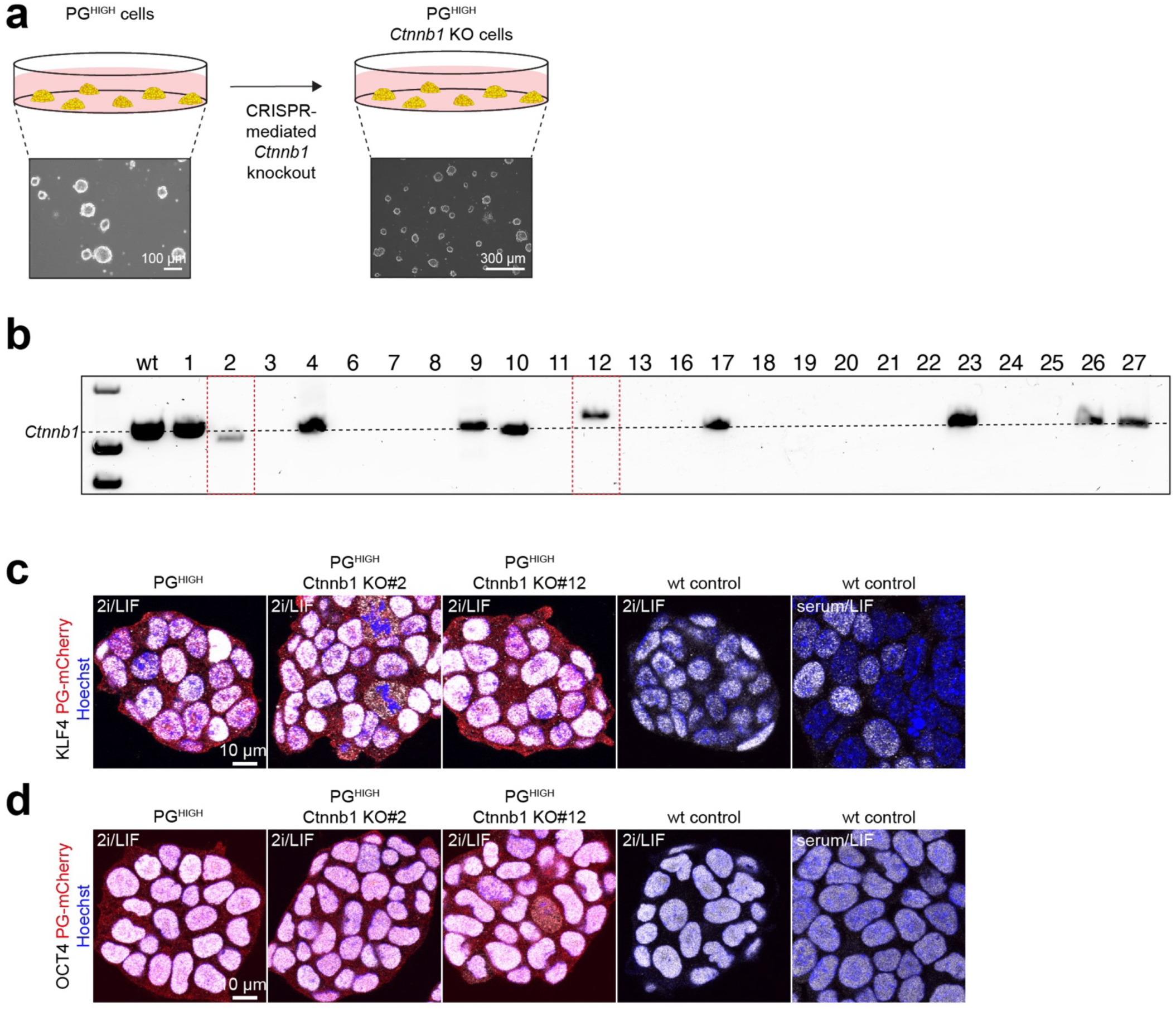
Plakoglobin sustains pluripotency independently of β-catenin. **a**, CRISPR-mediated knockout of *Ctnnb1* (β-catenin) in PG^HIGH^ cells. (see methods). **b**, Genomic knockout verification of *Ctnnb1* in PG^HIGH^ cells. *Ctnnb1* KO clone #2 and #12 are highlighted by the red dotted lines. **c-d**, Confocal immunofluorescence images of PG^HIGH^, PG^HIGH^ *Ctnnb1* KO (clone #2 and #12) and wild type control cells stained for KLF4 (naïve pluripotency marker) and OCT4 (general pluripotency marker). Scale bar: 10 µm.

## METHODS

### Data Availability Statement

The datasets generated during and/or analyzed during the current study are available in the GEO data repository, https://www.ncbi.nlm.nih.gov/geo/query/acc.cgi?acc=GSE197643.

Accession: GSE197643

### Cell culture

#### Mouse embryonic stem cells

The mouse embryonic stem cell lines E14 and Rex1GFPd2 (RGd2) were a kind gift from Austin Smith’s laboratory at the University of Cambridge. They were generally cultured in 2i/LIF medium^12^, which consist of N2B27 medium (1:1 DMEM/F-12 and Neurobasal media, N2 supplement^69^ (Cambridge Stem Cell Institute), B-27 supplement^70^ (gibco™), 0.11 % BSA fraction v (gibco™), 0.1 % sodium bicarbonate (gibco™), 12.5 µg/mL human insulin recombinat zinc (gibco™), 50 µM 2-mercaptoethanol (gibco™), 2 mM L-glutamine (gibco™), 1x penicillin-streptomycin (gibco™)) and is supplemented with 1 μM PD0325901 (Cambridge Stem Cell Institute), 3 μM CHIR99021 (Cambridge Stem Cell Institute), and 10 ng/mL leukemia inhibitor factor (Cambridge Stem Cell Institute). For a certain set of experiments, ESCs were cultured in serum/LIF medium (GMEM (Merck) was supplemented with 10% fetal calf serum (FCS) (Merck), 1x non-essential amino acids (NEAA) (Life Technologies), 1 mM sodium pyruvate (Life Technologies) and 1 mM L-Glutamine (Life Technologies)). Cells were maintained on gelatin-coated (0.1%) culture dishes in a 7% CO_2_ humidified incubator at 37 °C and medium was replaced every other day. Cells were passaged every 2 to 3 days at 80×10^4 cells/6-well (∼8,000 cells/cm^2^). To passage the cells or dissociate the cells into single cells for further experimental procedures the cells were treated as follows: the medium was aspirated, and cells were treated with Accutase^®^ at 37 °C until cells detached from the dish. Colonies were dissociated into single cells by gentle pipetting of the detached cells against the tissue culture dish. Afterwards, the Accutase^®^ was diluted with 10x volume of wash medium (DMEM-F12 + 5% BSA) and cell were centrifuged for 3 minutes at 300 RCF. Finally, the supernatant was discarded, cells were resuspended and replated in the appropriate pre-warmed medium.

### Embryo studies

#### Mouse embryo dissection and culture

All mice used were intercrosses of strain CD1 and obtained through natural mating. Embryonic staging assumed, that mating occurred at midnight. Hence, embryos were staged at E0.5 and noon of the following day. Pregnant females were killed on day 2.5, day 3.5 and day 4.5 post coitum (E2.5, E.3.5 and E4.5) by cervical dislocation. Oviduct and uterus were dissected and flushed with M2 medium (Merck) using Leica M165C stereo microscope. Post-implantation E5.5 and E6.5 embryos were carefully dissected from the implantation sites within the uterus and washed in M2 medium before fixation and subsequent processing.

#### Embryo injections for chimera generation

Oviduct and uterus were dissected and flushed with M2 medium (Merck) using Leica M165C stereo microscope. For (single) cell injections, ESCs grown in the appropriate conditions were dissociated by Accutase^®^ treatment for 5 minutes at 37 °C. Cells were then resuspended in washing buffer (9.5 mL, DMEM/F-12 + 5% BSA) and centrifuged for 3 minutes at 300 RCF. Subsequently, cells were resuspended in their appropriate medium (2i/LIF or serum/LIF) and stored on ice until injection. ESCs were loaded into a microinjection pipette and injected into the perivitelline space of 8-cell stage embryos via laser-generated perforation of the zona pellucida using the XYClone^®^ (Hamilton Thorne Biosciences). Subsequently, injected embryos were cultured in ORIGIO^®^ Blast^TM^ medium (Origio) at 37 °C and 7% CO_2_ for 48 hours, the equivalent of the E4.5 blastocysts.

#### Immunofluorescence staining of embryos

Initially, the zona pellucida was removed using acid Tyrode’s solution. Subsequently, embryos were fixed with 4% PFA for 20 minutes at room temperature. After one wash in PBS supplemented with 3 mg/mL polyvinylpyrolidone (PBS/PVP), embryos were permeabilized for ∼30 minutes in PBS/PVP with 0.25% Triton X-100. Afterwards, embryos were blocked in PBS containing 0.1% BSA, 0.01% Tween 20 and 2% donkey serum (blocking buffer). Primary antibodies were appropriately diluted in blocking buffer and incubated at 4 °C overnight. Subsequently, embryos were washed 3x in blocking buffer for at least 15 minutes each time before incubation with the secondary antibodies for 2 hours at room temperature and in the dark. Subsequently, embryos were washed in blocking buffer. Imagining was performed by placing the embryos in 2 μL droplets of water in a microscopy dish covered with mineral oil.

#### Human embryo culture, immunofluorescence staining and ethics statement

Use of human embryos in this work is approved by the Multi-Centre Research Ethics Committee, approval O4/MRE03/44 and approved by the Human Embryology & Fertilization Authority of the United Kingdom, research license R0178. Frozen embryos, surplus to requirement, were donated with informed consent by couples undergoing in vitro fertility treatment. Embryos were thawed using EmbryoThawTM Kit (EMF40_T, Fertipro) according to the manufacturer’s instructions. 4-cell stage embryos were cultured briefly in Cleave™ (Origio). Blastocysts were thawed directly into N2B27 medium (SCS-SF-NB-02, Stem Cell Sciences) and cultured to the desired stage in 50 μL drops of medium under mineral oil (MINOIL500, Fertipro) in a humidified atmosphere at 5%O_2_, 7% CO_2_ and 37 °C. Embryos were processed for immunohistochemistry by fixation in 4% paraformaldehyde in PBS for 15 minutes, rinsing in PBS containing 3 mg/mL polyvinylpyrolidone (PBS/PVP; P0930, Sigma), permeabilized in PBS/PVP containing 0.25% Triton X-100 (23,472-9, Sigma) for 30 minutes and blocked in buffer comprising PBS, 0.1% BSA, 0.01% Tween 20 (P1379, Sigma) and 2% donkey serum. Primary antibodies were diluted in blocking buffer and embryos incubated in antibody solution at 4 °C overnight. They were rinsed three times in blocking buffer for 15 minutes each, and incubated in secondary antibody solution for 1 hour at room temperature. Embryos were rinsed three times in blocking buffer, incubated briefly in increasing concentrations of Vectashield (H-1200, Vector Labs, Peterborough, UK) before mounting on glass slides in small drops of concentrated Vectashield with DAPI, and subsequently sealed with nail varnish.

### Bulk and scRNA-seq of mouse embryonic stem cells

#### Bulk RNA library preparations and sequencing

For each triplicate condition, a total of 1,000 cells were dispensed in single wells and processed using the Smart-seq2 protocol^71^. The amount of PCR cycles for the second-strand generation step were adjusted to 12 cycles to account for the increased input material. The unfragmented libraries were quality-controlled using a Bioanalyzer 2100 HS kit (Agilent). The libraries were then processed for tagmentation, using 1 ng of amplified cDNA as input, and 8 cycles were used for the dual-indexing PCR. The DNA molarity of the libraries after AMPureXP (Beckman Coulter) clean-up was assessed using a Bioanalyzer 2100 HS and a Qubit HS (Thermo Fischer) to estimate size distribution and DNA concentration respectively. Finally, the libraries were pooled at equimolar ratios and sequenced using a MiSeq v2 300 cycle kit (Illumina) (DNA Sequencing Facility, Dept. of Biochemistry).

#### Bulk RNA-seq data analysis

The fastq files were quality inspected using fastQC. The STAR aligner^72^ was used for mapping each demultiplexed paired-end file to a mm10 reference genome with a Gencode M12 gtf annotation file. Counting was achieved using featureCounts from the subread package^73^, and multi-mapped reads were discarded. Count tables were then imported in DESeq2^74^ for differential expression analysis between the triplicate conditions and Volcano plots were obtained using the EnhancedVolcano tool^75^. For GSEA analysis, we used the stat parameter provided from the DESeq2 gene expression analysis as an input for the WebGESTALT tool^76^.

#### inDrop single-cell RNA-seq library preparation

Cells were re-suspended at a final concentration of 120,000 cells/mL in PBS supplemented with 15% OptiPrep™ and libraries were prepared according to the inDrop protocol^77^ using the v3 barcoding configuration^78^. Briefly, each cell sample was encapsulated in water-in-oil emulsions containing a barcoded polyacrylamide microgel, a lysis buffer and reverse transcriptase mix. Each sample was collected in a single fraction containing approximately a 1,000 cells and libraries were processed according to the protocol. The bead-bound barcode was released from the hydrogel using UV exposure and the droplets were incubated at 50 °C for two hours followed by 70 °C for 20 minutes. The libraries were then processed for second-strand synthesis and amplification using *in vitro* transcription. After reverse-transcription, the libraries were amplified using limited-cycle indexing PCR. The final libraries were quality controlled using a BioAnalyzer HS kit (Agilent). Samples were pooled at equimolar ratios using both the BioAnalyzer HS and Qubit HS (ThermoFischer) metrics for size distribution and DNA concentration quantification. The libraries were sequenced on a Nextseq 75-cycle High Output kit (Illumina) in stand-alone mode using a 5% PhiX spike-in as an internal control.

#### Single-cell RNA-seq data analysis

The BCL files were converted to fastq files using the bcl2fastq script from Illumina. The read files were quality controlled using fastQC and de-multiplexed using Pheniqs^79^. The de-multiplexed files were used as an input for the zUMIs pipeline^80^ and mapped to a mouse GRCm38 reference genome with GRCm38.99 gtf annotation. The obtained aggregated intronic and exonic count matrices were then processed with the Scanpy tool^81^. Cells that had a low (<1%) or high fraction (>12%) of counts mapped to mitochondrial genes were excluded. Additionally, cells were filtered on the detected number of genes (between 800 and 3,000) and transcript counts (between 1,500 and 50,000). The adata objects were then normalized, and transcript counts were regressed out. After scaling and principal component analysis, a dimensional reduction plot was obtained by computing a two-dimensional Uniform Manifold Approximation and Projection (UMAP) was achieved using the scanpy.tl.umap() function. Clustering was achieved using the leiden algorithm (sc.tl.leiden()). Dynamical modelling of RNA velocity was performed with the scVelo tool^82^ using the velocyto loom files^83^ obtained via the zUMIs pipeline as an input. Latent times were computed using the scVelo package.

### Microscopy

#### Confocal microscopy

Imaging was performed with inverted Leica TCS SP8 confocal microscopes using 40x/1.30 and 20x/0.75 objectives. Fluorophores were excited with a 405 nm, a 488 nm, a 552 nm and a 638 nm laser. Raw data were analyzed with the open source software ImageJ/Fiji.

#### Atomic force microscopy

AFM force measurements were performed on a JPK CellHesion 200 AFM (Bruker) using an Arrow TL1 cantilevers (NanoWorld) onto which a 37 mm polystyrene bead (microParticles) had been glued. For sample preparation low melting agarose (lonza) was mixed with PBS (1.5% w/v), dissolved at 70 °C, and subsequently cooled down to 37 °C. 2 mL were poured into TTP™ dishes (P/N 93040), cooled down in the fridge overnight and PBS was either added directly before the measurement or 1.5 hours before. For the measurements set points of up to 60 nN were chosen and the speed of the Z scanner was 5 mm/s. Each gel was measured in 16 locations. The resulting force-distance curves were then analyzed in the JPK Data Processing software, where the Hertz Model was applied to calculate the Young’s Modulus, assuming Poisson’s ratio to be 0.5. Care was taken that the indentation remained below one third of the bead radius.

### Microfluidics

#### Fabrication of master molds

The microfluidic chips were designed with the computer-aided design (CAD) software (DraftSight) and were produced using well-established soft-lithography protocols^84^. Briefly, chip designs were cut out on a photo mask which was then used to fabricate of a silicon wafer master mold in a clean room.

#### Microfluidic chip production

The silicon master molds were placed in a petri dish and filled with SYLGARD™ 184 silicone elastomer base and curing agent (Merck) in a 10:1 (w/w) ratio. After degassing, the PDMS was polymerized at 65 °C overnight and carefully removed from the master mold. The in- and outlets for the chip were generated with a 1 mm biopsy puncher (pmf medical). After thorough cleaning with 2-propanol and compressed air, the PDMS and microscope slides were placed into a low-pressure oxygen plasma generator. A vacuum (0 mbar) was generated and oxygen plasma introduced for 12 seconds. The PDMS was pressed onto the microscope slides to form covalent bonds, the channels where coated/treated with 1% (v/v) Trichloro(1H,1H,2H,2H-perfluorooctyl)silane (Merck) in HFE-7500 (Fluorochem) and the chips were placed on a 50 °C hot plate for 30 minutes. Until use, chips were stored in a sterile container.

#### Encapsulation

Low melting-point agarose (Lonza) was mixed with PBS, heated up to 70 °C to form a 3% (w/v) solution and kept at 37 °C until the encapsulation. A single cell suspension was mixed gently but thoroughly with the agarose solution (1:1 ratio) and carefully aspirated into a polyethene tubing (Portex^®^) connected to a glass syringe (SEG Analytical Science). Another syringe was filled with HFE-7500 oil (Fluorochem) and Pico-Surf surfactant (Sphere Fluidics) with the final concentration of 0.3%. The Syringes were connected to the microfluidic chip and automated pumps (CETONI) injected both the oil-surfactant solution and the agarose-cell suspension through different channels. Agarose droplets were formed and collected on ice to allow polymerization of the agarose. Using liquid-liquid extraction the microgels were demulsified with 1H,1H,2H,2H-perfluoro-1-octanol (AlfaAesar) and culture medium.

#### Cell recovery from microgels

For the recovery of ESCs from agarose microgel, the cell culture medium including the microgels was collected in a tube and gently centrifuged at 200 RCF for 3 minutes. The medium was aspirated until only 1 mL was remaining and 2 µL Agarase (0.5 U/µL, Thermo Fischer) was added and gently mixed. They were incubated at 37 °C until the microgels were dissolved and the cell colonies released (∼5-10 minutes). The cell colonies were then either used for further experiments, after the transfer into new medium, or they were pelleted at 300 RCF for 3 minutes and treated with Accutase^®^ to generate single cell suspension for example for flow cytometry analysis.

### Molecular biology

#### Cloning of the *Jup*-*T2A-mCherry* plasmid

The transposon for the genomic integration of *Jup* using the PiggyBac system was constructed as follows. The *Jup* cDNA was recovered from the bulk RNA-seq experiment cDNA pool and amplified for downstream vector assembly. The PCR-amplified *Jup* cDNA (primer pair: F_Jup-cDNA and R_Jup-cDNA) was inserted into the previously described pPB-CAG vector^85^ together with an mCherry gene via Gibson assembly. For bi-cistronic expression under the control of the CAG promoter, mCherry was linked to Jup via the ribosome-skipping 2A sequence of *thosea asigna* virus (T2A). The final *PB-CAG-Jup-T2A-mCherry* construct was validated by Sanger sequencing (DNA Sequencing Facility, Dept. of Biochemistry).

#### Generation of Plakoglobin overexpressing RGd2 ESCs

To generate the *Jup* overexpressing cell line, the PiggyBac transposon system^85^ was used. Briefly, 10 µL Lipofectamine 2000 were mixed with 300 µL 2i/LIF medium. Separately, 1.2 µL *PB_Jup-T2A-mCherry* construct (700 ng/mL, 0.8 µg) with 1.6 µL PBase (500 ng/mL, 0.75 µg) and 150 µL 2i/LIF medium. Then, 150 µL diluted DNA were mixed with 300 µL diluted Lipofectamine 2000 and incubated for 5 minutes at room temperature. Subsequently, ∼5×10^5^ cells were resuspended in 450 µL of DNA/Lipofectamine 2000 and incubated for 10 minutes at 37 °C. Then, cells were plated and stably transfected cells FACS-sorted based on their mCherry signal.

#### *Ctnnb1* knock-out via CRISPR Cas9

β-catenin knockout cells were generated via transient transfection with a CRISPR/Cas9 *Ctnnb1* KO plasmid according to the manufacturer’s instructions (Santa Cruz, sc-419477). For knock-out validation, genomic DNA of grown clones was harvested using QuickExtract™ DNA Extraction Solution (*Lucigen*) following the manufacturer’s instructions. The solution was directly used as a template for PCR amplification using Q5^®^ High-Fidelity 2X Master Mix (*NEB*) (primer pair: F_Ctnnb1_Exon3to6 and R_Ctnnb1_Exon3to6). PCR products were resolved by agarose gel electrophoresis and relevant bands extracted and purified for Sanger sequencing (DNA Sequencing Facility, Dept. of Biochemistry) to identify genome edit at targeted sites.

### Cell analysis

#### Hanging drop culture

A single cell suspension with either 333 cells/mL or 33 cells/mL was generated in serial dilution with 2i/LIF medium with prior Accutase^®^ treatment of ESCs. Subsequently, 30 µL drops (containing ∼10 cells or ∼1 cell) were distributed onto the inside of a 15 cm petri dish and the lid was placed onto the PBS filled dish, so that the drops were hanging upside down in the lid without touching the PBS.

#### Clonogenicity assay

mESCs cultured in 2i/LIF or serum/LIF were either cultured on TCP or encapsulated into agarose microgels. After 48 hours the microgels were dissolved with Agarase (Thermo Fisher) and single cell suspensions were generated with Accutase^®^, both from cells on TCP and microgels. 1,000 cells were seeded in 2i/LIF on gelatin-coated 6-wells (cell density of ∼10^4^ cells/cm^2^). Surviving colonies were counted after 48 hours.

#### Flow cytometry and cell sorting

The fluorescent cell reporters Rex1-GFPd2 and PG-mCherry were analyzed with either the LSRFortessa™ (BD Biosciences) or the Attune™ NxT (Thermo Fisher) flow cytometer. Single cell suspensions were generated by Accutase^®^ treatment. The cells were washed with DMEM-F12 + 5% BSA, resuspended in PBS or medium and kept on ice until the measurement. The data were analyzed with the software FlowJo (BD Biosciences).

#### Fluorescence activated cell sorting

Bulk and single cell fluorescence activated cell sorting (FACS) was performed at the Cambridge Stem Cell Institute Flow Facility on a MoFlow XDP cell sorter (Beckmann Coulter) and the Flow Cytometry School of the biological sciences on the BD FACSAria™ IIu Cell Sorter (BD Biosciences). The cells were treated with Accuatse^®^, washed with DMEM-F12 + 5% BSA, resuspended in cell media and stored on ice until FACS. Bulk sorts were collected in a collection tube and washed after the sorting before culturing the cells in a gelatin-coated 6-well plate. Single cells were directly sorted into gelatin-coated 96-well plates.

#### Immunofluorescence staining of cells

Cells were fixed for 10 minutes with 4% formaldehyde solution (Thermo Fischer) in PBS. After washing with PBS, cells were permeabilized for 15 minutes with 0.25% Triton X-100 in PBS. Subsequently, cells were blocked for at least 2 hours with 3% BSA in PBS blocking solution. Primary antibody incubation was performed at desired concentration at 4 °C, overnight. Cells were then washed with blocking buffer and incubated with the secondary antibody for 2 hours at room temperature. Afterwards cells were washed with blocking buffer and stained with 1 µg/mL Hoechst.

## References

1. Rheinlaender, J. et al. Cortical cell stiffness is independent of substrate mechanics. Nat Mater 19, 1019–1025 (2020).

2. De Belly, H. et al. Membrane Tension Gates ERK-Mediated Regulation of Pluripotent Cell Fate. Cell Stem Cell 28, 273–284 e276 (2021).

3. Bergert, M. et al. Cell Surface Mechanics Gate Embryonic Stem Cell Differentiation. Cell Stem Cell 28, 209–216 e204 (2021).

4. Li, Y.W. et al. Volumetric Compression Induces Intracellular Crowding to Control Intestinal Organoid Growth via Wnt/beta-Catenin Signaling (vol 28, pg 1, 2021). Cell Stem Cell 28, 170–172 (2021).

5. Chan, C.J., Heisenberg, C.P. & Hiiragi, T. Coordination of Morphogenesis and Cell-Fate Specification in Development. Curr Biol 27, R1024–R1035 (2017).

6. Heisenberg, C.P. & Bellaiche, Y. Forces in tissue morphogenesis and patterning. Cell 153, 948–962 (2013).

7. Evans, M.J. & Kaufman, M.H. Establishment in culture of pluripotential cells from mouse embryos. Nature 292, 154–156 (1981).

8. Martin, G.R. Isolation of a pluripotent cell line from early mouse embryos cultured in medium conditioned by teratocarcinoma stem cells. Proc Natl Acad Sci U S A 78, 7634–7638 (1981).

9. Boroviak, T., Loos, R., Bertone, P., Smith, A. & Nichols, J. The ability of inner-cell-mass cells to self-renew as embryonic stem cells is acquired following epiblast specification. Nat Cell Biol 16, 516–528 (2014).

10. Boroviak, T. et al. Lineage-Specific Profiling Delineates the Emergence and Progression of Naive Pluripotency in Mammalian Embryogenesis. Dev Cell 35, 366–382 (2015).

11. Bradley, A., Evans, M., Kaufman, M.H. & Robertson, E. Formation of germ-line chimaeras from embryo-derived teratocarcinoma cell lines. Nature 309, 255–256 (1984).

12. Ying, Q.L. et al. The ground state of embryonic stem cell self-renewal. Nature 453, 519–523 (2008).

13. Alexandrova, S. et al. Selection and dynamics of embryonic stem cell integration into early mouse embryos. Development 143, 24–34 (2016).

14. Burdon, T., Stracey, C., Chambers, I., Nichols, J. & Smith, A. Suppression of SHP-2 and ERK signalling promotes self-renewal of mouse embryonic stem cells. Dev Biol 210, 30–43 (1999).

15. Kunath, T. et al. FGF stimulation of the Erk1/2 signalling cascade triggers transition of pluripotent embryonic stem cells from self-renewal to lineage commitment. Development 134, 2895–2902 (2007).

16. Wray, J. et al. Inhibition of glycogen synthase kinase-3 alleviates Tcf3 repression of the pluripotency network and increases embryonic stem cell resistance to differentiation. Nat Cell Biol 13, 838–845 (2011).

17. Lyashenko, N. et al. Differential requirement for the dual functions of beta-catenin in embryonic stem cell self-renewal and germ layer formation. Nat Cell Biol 13, 753–761 (2011).

18. Dunn, S.J., Martello, G., Yordanov, B., Emmott, S. & Smith, A.G. Defining an essential transcription factor program for naive pluripotency. Science 344, 1156–1160 (2014).

19. Martello, G., Bertone, P. & Smith, A. Identification of the missing pluripotency mediator downstream of leukaemia inhibitory factor. Embo J 32, 2561–2574 (2013).

20. Wray, J., Kalkan, T. & Smith, A.G. The ground state of pluripotency. Biochem Soc Trans 38, 1027–1032 (2010).

21. Chowdhury, F. et al. Soft substrates promote homogeneous self-renewal of embryonic stem cells via downregulating cell-matrix tractions. PLoS One 5, e15655 (2010).

22. Chowdhury, F. et al. Material properties of the cell dictate stress-induced spreading and differentiation in embryonic stem cells. Nat Mater 9, 82–88 (2010).

23. Przybyla, L., Lakins, J.N. & Weaver, V.M. Tissue Mechanics Orchestrate Wnt-Dependent Human Embryonic Stem Cell Differentiation. Cell Stem Cell 19, 462–475 (2016).

24. Przybyla, L.M., Theunissen, T.W., Jaenisch, R. & Voldman, J. Matrix remodeling maintains embryonic stem cell self-renewal by activating Stat3. Stem Cells 31, 1097–1106 (2013).

25. Labouesse, C. et al. StemBond hydrogels control the mechanical microenvironment for pluripotent stem cells. Nat Commun 12, 6132 (2021).

26. Engler, A.J., Sen, S., Sweeney, H.L. & Discher, D.E. Matrix elasticity directs stem cell lineage specification. Cell 126, 677–689 (2006).

27. Gudipaty, S.A. et al. Mechanical stretch triggers rapid epithelial cell division through Piezo1. Nature 543, 118–121 (2017).

28. Segel, M. et al. Niche stiffness underlies the ageing of central nervous system progenitor cells. Nature 573, 130–134 (2019).

29. Verstreken, C.M., Labouesse, C., Agley, C.C. & Chalut, K.J. Embryonic stem cells become mechanoresponsive upon exit from ground state of pluripotency. Open Biol 9, 180203 (2019).

30. Li, Y. et al. Compression-induced dedifferentiation of adipocytes promotes tumor progression. Sci Adv 6, eaax5611 (2020).

31. Wang, X. et al. Characterizing Inner Pressure and Stiffness of Trophoblast and Inner Cell Mass of Blastocysts. Biophys J 115, 2443–2450 (2018).

32. Chan, C.J. et al. Hydraulic control of mammalian embryo size and cell fate. Nature 571, 112–116 (2019).

33. Kalkan, T. et al. Tracking the embryonic stem cell transition from ground state pluripotency. Development 144, 1221–1234 (2017).

34. Kleine-Bruggeney, H. et al. Long-Term Perfusion Culture of Monoclonal Embryonic Stem Cells in 3D Hydrogel Beads for Continuous Optical Analysis of Differentiation. Small 15 (2019).

35. Mulas, C. et al. Microfluidic platform for 3D cell culture with live imaging and clone retrieval. Lab Chip (2020).

36. Schindler, M. et al. Agarose microgel culture delineates lumenogenesis in naive and primed human pluripotent stem cells. Stem Cell Reports 16, 1347–1362 (2021).

37. Marks, H. et al. The transcriptional and epigenomic foundations of ground state pluripotency. Cell 149, 590–604 (2012).

38. Peifer, M., Mccrea, P.D., Green, K.J., Wieschaus, E. & Gumbiner, B.M. The Vertebrate Adhesive Junction Proteins Beta-Catenin and Plakoglobin and the Drosophila Segment Polarity Gene Armadillo Form a Multigene Family with Similar Properties. J Cell Biol 118, 681–691 (1992).

39. Salomon, D. et al. Regulation of beta-catenin levels and localization by overexpression of plakoglobin and inhibition of the ubiquitin-proteasome system. J Cell Biol 139, 1325–1335 (1997).

40. Klein, A.M. et al. Droplet barcoding for single-cell transcriptomics applied to embryonic stem cells. Cell 161, 1187–1201 (2015).

41. La Manno, G. et al. RNA velocity of single cells. Nature 560, 494–498 (2018).

42. Bergen, V., Lange, M., Peidli, S., Wolf, F.A. & Theis, F.J. Generalizing RNA velocity to transient cell states through dynamical modeling. Nat Biotechnol 38, 1408–1414 (2020).

43. Buecker, C. et al. Reorganization of enhancer patterns in transition from naive to primed pluripotency. Cell Stem Cell 14, 838–853 (2014).

44. Kolodziejczyk, A.A. et al. Single Cell RNA-Sequencing of Pluripotent States Unlocks Modular Transcriptional Variation. Cell Stem Cell 17, 471–485 (2015).

45. Shyh-Chang, N. et al. Influence of threonine metabolism on S-adenosylmethionine and histone methylation. Science 339, 222–226 (2013).

46. Tischler, J. et al. Metabolic regulation of pluripotency and germ cell fate through alpha-ketoglutarate. Embo J 38 (2019).

47. Huang, S.M. et al. Tankyrase inhibition stabilizes axin and antagonizes Wnt signalling. Nature 461, 614–620 (2009).

48. Guilak, F. et al. Control of stem cell fate by physical interactions with the extracellular matrix. Cell Stem Cell 5, 17–26 (2009).

49. Crowder, S.W., Leonardo, V., Whittaker, T., Papathanasiou, P. & Stevens, M.M. Material Cues as Potent Regulators of Epigenetics and Stem Cell Function. Cell Stem Cell 18, 39–52 (2016).

50. Zujur, D. et al. Three-dimensional system enabling the maintenance and directed differentiation of pluripotent stem cells under defined conditions. Sci Adv 3, e1602875 (2017).

51. Caiazzo, M. et al. Defined three-dimensional microenvironments boost induction of pluripotency. Nat Mater 15, 344–352 (2016).

52. Cowin, P., Kapprell, H.P., Franke, W.W., Tamkun, J. & Hynes, R.O. Plakoglobin - a Protein Common to Different Kinds of Intercellular Adhering Junctions. Cell 46, 1063–1073 (1986).

53. Bierkamp, C., McLaughlin, K.J., Schwarz, H., Huber, O. & Kemler, R. Embryonic heart and skin defects in mice lacking plakoglobin. Dev Biol 180, 780–785 (1996).

54. Fernandez-Sanchez, M.E. et al. Mechanical induction of the tumorigenic beta-catenin pathway by tumour growth pressure. Nature 523, 92–95 (2015).

55. Muncie, J.M. et al. Mechanical Tension Promotes Formation of Gastrulation-like Nodes and Patterns Mesoderm Specification in Human Embryonic Stem Cells. Dev Cell 55, 679–694 e611 (2020).

56. Brunet, T. et al. Evolutionary conservation of early mesoderm specification by mechanotransduction in Bilateria. Nat Commun 4, 2821 (2013).

57. Weber, G.F., Bjerke, M.A. & DeSimone, D.W. A mechanoresponsive cadherin-keratin complex directs polarized protrusive behavior and collective cell migration. Dev Cell 22, 104–115 (2012).

58. Ohsugi, M. et al. Expression and cell membrane localization of catenins during mouse preimplantation development. Dev Dyn 206, 391–402 (1996).

59. Mahendram, S. et al. Ectopic gamma-catenin expression partially mimics the effects of stabilized beta-catenin on embryonic stem cell differentiation. PLoS One 8, e65320 (2013).

60. Shimizu, M., Fukunaga, Y., Ikenouchi, J. & Nagafuchi, A. Defining the roles of beta-catenin and plakoglobin in LEF/T-cell factor-dependent transcription using beta-catenin/plakoglobin-null F9 cells. Mol Cell Biol 28, 825–835 (2008).

61. Huelsken, J. et al. Requirement for beta-catenin in anterior-posterior axis formation in mice. J Cell Biol 148, 567–578 (2000).

62. Haegel, H. et al. Lack of beta-catenin affects mouse development at gastrulation. Development 121, 3529–3537 (1995).

63. Miller, J.R. & Moon, R.T. Analysis of the signaling activities of localization mutants of beta-catenin during axis specification in Xenopus. J Cell Biol 139, 229–243 (1997).

64. Yi, F. et al. Opposing effects of Tcf3 and Tcf1 control Wnt stimulation of embryonic stem cell self-renewal. Nat Cell Biol 13, 762–770 (2011).

65. Shy, B.R. et al. Regulation of Tcf7l1 DNA binding and protein stability as principal mechanisms of Wnt/beta-catenin signaling. Cell Rep 4, 1–9 (2013).

66. Valet, M., Siggia, E.D. & Brivanlou, A.H. Mechanical regulation of early vertebrate embryogenesis. Nat Rev Mol Cell Biol (2021).

67. Broders-Bondon, F., Nguyen Ho-Bouldoires, T.H., Fernandez-Sanchez, M.E. & Farge, E. Mechanotransduction in tumor progression: The dark side of the force. J Cell Biol 217, 1571–1587 (2018).

## References

68. Mohammed, H. et al. Single-Cell Landscape of Transcriptional Heterogeneity and Cell Fate Decisions during Mouse Early Gastrulation. Cell Rep 20, 1215–1228 (2017).

69. Bottenstein, J.E. & Harvey, A.L. Cell Culture in the Neurosciences. Plenum Press: New York and London., 3 (1985).

70. Brewer, G.J., Torricelli, J.R., Evege, E.K. & Price, P.J. Optimized survival of hippocampal neurons in B27-supplemented Neurobasal, a new serum-free medium combination. J Neurosci Res 35, 567–576 (1993).

71. Picelli, S. et al. Full-length RNA-seq from single cells using Smart-seq2. Nature Protocols 9, 171–181 (2014).

72. Dobin, A. et al. STAR: ultrafast universal RNA-seq aligner. Bioinformatics 29, 15–21 (2012).

73. Liao, Y., Smyth, G.K. & Shi, W. featureCounts: an efficient general purpose program for assigning sequence reads to genomic features. Bioinformatics 30, 923–930 (2013).

74. Love, M., Ahlmann-Eltze, C., Forbes, K., Anders, S. & Huber, W. Differential gene expression analysis based on the negative binomial distribution. Bioconductor.

75. Blighe, K., Rana, S. & Lewis, M. EnhancedVolcano: Publication-ready volcano plots with enhanced colouring and labeling. R package version 1.12.0 (2021).

76. Liao, Y., Wang, J., Jaehnig, E.J., Shi, Z. & Zhang, B. WebGestalt 2019: gene set analysis toolkit with revamped UIs and APIs. Nucleic Acids Research 47, W199–W205 (2019).

77. Zilionis, R. et al. Single-cell barcoding and sequencing using droplet microfluidics. Nature Protocols 12, 44–73 (2017).

78. Briggs, J.A. et al. The dynamics of gene expression in vertebrate embryogenesis at single-cell resolution. Science 360, eaar5780 (2018).

79. Galanti, L., Shasha, D. & Gunsalus, K.C. Pheniqs 2.0: accurate, high-performance Bayesian decoding and confidence estimation for combinatorial barcode indexing. BMC Bioinformatics 22, 359 (2021).

80. Parekh, S., Ziegenhain, C., Vieth, B., Enard, W. & Hellmann, I. zUMIs - A fast and flexible pipeline to process RNA sequencing data with UMIs. GigaScience 7 (2018).

81. Wolf, F.A., Angerer, P. & Theis, F.J. SCANPY: large-scale single-cell gene expression data analysis. Genome Biology 19, 15 (2018).

82. Bergen, V., Lange, M., Peidli, S., Wolf, F.A. & Theis, F.J. Generalizing RNA velocity to transient cell states through dynamical modeling. Nature Biotechnology 38, 1408–1414 (2020).

83. La Manno, G. et al. RNA velocity of single cells. Nature 560, 494–498 (2018).

84. Xia, Y. & Whitesides, G.M. Soft Lithography. Angew Chem Int Ed Engl 37, 550–575 (1998).

85. Yusa, K., Rad, R., Takeda, J. & Bradley, A. Generation of transgene-free induced pluripotent mouse stem cells by the piggyBac transposon. Nat Methods 6, 363–369 (2009).

