## Supplemental Methods Table for "Plakoglobin is a mechanoresponsive regulator of naïve pluripotency"

#### Material Table

| Antibodies | Source | Identifier |
|---|---|---|
| α-E-catenin | Cell Signaling Technology | #3236 |
| Lamin B1 | abcam | ab16048 |
| Oct-3/4 (C-10) | Santa Cruz Biotechnology | sc-5279 |
| Catenin δ-1 | Cell Signaling Technology | #59854 |
| E-cadherin | Thermo Fisher Scientific | # 13-1900 |
| ERR beta/NR3B2 | R&D Systems | PP-H6705-00 |
| SOX17 | R&D Systems | AF1924 |
| SOX2 | R&D Systems | MAB2018 |
| KLF4 | R&D Systems | AF3158 |
| Nanog | Thermo Fisher Scientific | # 14-5761-80 |
| Rabbit mAB β-Catenin | Cell Signaling Technology | #8480 |
| T Brachury | R&D Systems | AF2085 |
| γ-Catenin Antibody | Cell Signaling Technology | #2309 |

| Assays | Source | Identifier |
|---|---|---|
| AMPure XP | Beckman Coulter | A63880 |
| BioAnalyzer High Sensitivity DNA Analysis | Agilent | 5067-4626 |
| MiSeq Reagent Kit V2 (300 cycle kit) | Illumina | MS-102-2002 |
| NextSeq 500/550 High Output Kit v2.5 (75 c.) | Illumina | 20024906 |
| Qubit 1X dsDNA Assay Kit, high sensitivity | Thermo Fisher Scientific | Q33230 |

| Chemicals | Source | Identifier |
|---|---|---|
| 16% Formaldehyde solution (w/v) | Thermo Fisher Scientific | 28908 |
| 1H,1H,2H,2H-Perfluorooctanol (97 %) | AlfaAesar | B20156 |
| 2-Mercaptoethanol | gibco | 31350-010 |
| Accutase | Merck | A6964 |
| Agarase | Thermo Fisher Scientific | EO0461 |
| B-27 Supplement | gibco | 17504-044 |
| Bovine Albumin Fraction V (7.5% solution) | Thermo Fisher Scientific | 15260037 |
| BSA Fraction V (7.5%) | gibco | 15260-037 |
| CHIR99021 | Cambridge Stem Cell Institute | N/A |
| DMEM/F-12 | gibco | 21331-020 |
| FCS | MRC Cambridge Stem Cell Institute | N/A |
| GMEM | Sigma | G5154 |
| HFE-7500 3M Novec Engineered fluid | Fluorochem Ltd | 51243 |
| Insulin, Human Recombinant Zinc | gibco | 12585-014 |
| L-glutamine | gibco | 25030-081 |
| Leukaemia inhibitor factor | Cambridge Stem Cell Institute | N/A |
| Lipofectamine 2000 | Thermo Fisher Scientific | 11668030 |
| M2 medium | Merck | M7167 |
| MEM Non-Essential Amino Acids Solution | gibco | 11140-035 |
| N2 | Cambridge Stem Cell Institute | |
| Neurobasal | gibco | 21103-049 |
| OptiPrep | Stemcell Technologies | 7820 |
| ORIGIO Sequential Blast | Origio | 83050010A |
| Paraformaldehyde 16 % | AlfaAesar | 43368 |
| PD0325901 | Cambridge Stem Cell Institute | N/A |
| Penicillin-Streptomycin | gibco | 15140-122 |
| Pico-Surf | Sphere Fluidics | C021 |
| Q5® High-Fidelity 2X Master Mix | New England BioLabs | M0492S |
| QuickExtract DNA Extraction Solution | Lucigen | QE09050 |
| SeaPrep Agarose | Lonza | 50302 |
| Sodium Bicarbonate | gibco | 25080-094 |
| Sodium pyruvate | gibco | 11360-070 |
| SYLGARD 184 Silicone Elastomer Kit | Merck | 761036 |

| | | |
|---|---|---|
| Trichloro-(1H,1H,2H,2H-perfluoro-octyl)silane | Merck | 448931 |
| **Eukaryotic cell lines** | **Source** | **Identifier** |
| E14TG2a | Prof. Austin Smith's lab | N/A |
| PG-OE | This study | N/A |
| PGHIGH#1 | This study | N/A |
| PGHIGH#2 | This study | N/A |
| PGHIGH#2 Ctnnb1 KO #12 | This study | N/A |
| PGHIGH#2 Ctnnb1 KO #2 | This study | N/A |
| PGHIGH#3 | This study | N/A |
| PGLOW#1 | This study | N/A |
| PGLOW#2 | This study | N/A |
| PGLOW#3 | This study | N/A |
| RGd2 | [1] | N/A |
| **Molecular biology** | **Source** | **Identifier** |
| F_Ctnnb1-Exon3to6: CAACCCTGAGGAAGAAGATGTTGACACC | This study | N/A |
| F_Jup-cDNA: gtctcatcattttggcaaagaattcccATGGAG GTGATGAACCTTATTGAGCAG | This study | N/A |
| gRNA1: ATGAGCAGCGTCAAACTGCG | Santa Cruz | sc-419477 |
| gRNA2: AGCTACTTGCTCTTGCGTGA | Santa Cruz | sc-419477 |
| gRNA3: AAAATGGCAGTGCGCCTAGC | Santa Cruz | sc-419477 |
| R_Ctnnb1-Exon3to6: GCTAAGATCTGAAGGCAGTCTGTTGTAATAGCC | This study | N/A |
| R_Jup-cDNA: agcagacttcctctgccctcGGCCAGCATGTGGTCTGC | This study | N/A |
| **Software and code** | **Source** | |
| bcl2fastq Conversion | Illumina | |
| Computer-aided design software | DraftSight | |
| Data Processing software | JPK | |
| DESeq2 | [2] | |
| EnhancedVolcano tool | [3] | |
| featureCounts | [4] | |
| Fiji | [5] | |
| FlowJo | FlowJo LLC | |
| Illustrator | Adobe Systems | |
| Pheniqs 2.0 | [6] | |
| Python | Python Software Foundation | |
| R | R Core Team | |
| Scanpy tool | [7] | |
| scVelo | [8] | |
| STAR: ultrafast universal RNA-seq aligner | [9] | |
| WebGestalt | [10] | |

### References


1. Wray, J. *et al.* Inhibition of glycogen synthase kinase-3 alleviates Tcf3 repression of the pluripotency network and increases embryonic stem cell resistance to differentiation. *Nature Cell Biology* **13**, 838-845 (2011).
2. Love, M., Ahlmann-Eltze, C., Forbes, K., Anders, S. & Huber, W. Differential gene expression analysis based on the negative binomial distribution. *Bioconductor*.
3. Blighe, K., Rana, S. & Lewis, M. EnhancedVolcano: Publication-ready volcano plots with enhanced colouring and labeling. *R package version 1.12.0* (2021).
4. Liao, Y., Smyth, G.K. & Shi, W. featureCounts: an efficient general purpose program for assigning sequence reads to genomic features. *Bioinformatics* **30**, 923-930 (2013).
5. Schindelin, J. *et al.* Fiji: an open-source platform for biological-image analysis. *Nature Methods* **9**, 676-682 (2012).
6. Galanti, L., Shasha, D. & Gunsalus, K.C. Pheniqs 2.0: accurate, high-performance Bayesian decoding and confidence estimation for combinatorial barcode indexing. *BMC Bioinformatics* **22**, 359 (2021).
7. Wolf, F.A., Angerer, P. & Theis, F.J. SCANPY: large-scale single-cell gene expression data analysis. *Genome Biology* **19**, 15 (2018).
8. Bergen, V., Lange, M., Peidli, S., Wolf, F.A. & Theis, F.J. Generalizing RNA velocity to transient cell states through dynamical modeling. *Nature Biotechnology* **38**, 1408-1414 (2020).
9. Dobin, A. *et al.* STAR: ultrafast universal RNA-seq aligner. *Bioinformatics* **29**, 15-21 (2012).
10. Liao, Y., Wang, J., Jaehnig, E.J., Shi, Z. & Zhang, B. WebGestalt 2019: gene set analysis toolkit with revamped UIs and APIs. *Nucleic Acids Research* **47**, W199-W205 (2019).
